# Leviathan: A fast, memory-efficient, and scalable taxonomic and pathway profiler for (pan)genome-resolved metagenomics and metatranscriptomics

**DOI:** 10.1101/2025.07.14.664802

**Authors:** Josh L. Espinoza, Allan Phillips, Chris L. Dupont

**Affiliations:** NewAtlantis Labs; J. Craig Venter Institute; Ocean BioMetrics; University of California, San Diego

## Abstract

Functional profiling of meta-omics is essential for understanding microbial communities, yet support for custom genome-resolved reference databases is limited. We introduce *Leviathan* for integrated taxonomic and functional profiling at both genome and pangenome resolution. *Leviathan* combines *Sylph* for ultrafast alignment-free taxonomic profiling with *Salmon* for pseudo-alignment-based read quantification in DNA-space against (pan)genome-resolved gene catalogs, producing dual metrics per (pan)genome: pathway abundance and graph-based pathway coverage. Benchmarking alignment backends on synthetic metagenomes, we show that DNA-space pseudo-alignments retain competitive (pan)genome-level classification performance compared to traditional alignment, reducing computational runtime and memory requirements, while translated searches in protein-space lose classification resolution from ambiguous mapping events. *Leviathan’s* utility is demonstrated through two case studies: a marine plastisphere metagenomics dataset analyzing metabolic shifts between early and mature biofilm communities and a dental caries metatranscriptomics dataset where co-expression network analysis identified organism-specific transcriptional patterns diagnostic of health and disease states. *Leviathan* is available at https://github.com/jolespin/leviathan.

**Importance:** Understanding what microbes can do, not just which ones are present, is central to translating microbiome research into actionable insight. Existing functional profiling tools either rely on fixed reference databases or require complex multi-step pipelines when applied to custom genome collections, and none natively compute per-(pan)genome pathway abundance and graph-based pathway completeness in a single workflow. This limits the ability for researchers to directly compare functional profiles to tangential analyses on their specific genome catalogs. Leviathan addresses this gap with integrated taxonomic and functional profiling against user-defined (pan)genome-resolved references using pseudo-alignment, achieving competitive classification accuracy with substantially lower runtime and memory requirements compared to current methods. Native pangenome support enables routine quantification of metabolic potential and transcriptional activity at both genome and pangenome resolution, revealing functional variation across related strains that single-genome or community-level analyses obscure.

## Introduction

High-throughput sequencing has revolutionized microbiology, enabling deep insights into the composition and function of complex microbial communities through metagenomics and metatranscriptomics. While taxonomic profiling reveals “who is there” and “how abundant they are relative to each other”, functional profiling uncovers “what they can do” providing crucial information for understanding ecosystem dynamics, host-microbe interactions, and biogeochemical cycles.

Contextualizing genetic assets with sample metadata is essential for translating basic research into actionable insight with genome-resolved metagenomics as a powerful approach for building scalable science. Taxonomic profiling can be implemented using *k*-mer based methods (e.g., *Sylph* (1), *Kraken2* (2), *Ganon* (3)) requiring only a reference database of genomes or marker-based methods (*MetaPhlAn* (4), *mOTUs* (5)) requiring both reference genomes and taxa-specific marker sets. While the methodologies for taxonomic profiling are well established and scalable, the development of functional profiling methodologies has lagged. Accurate and comprehensive functional profiling from meta-omics datasets requires: 1) establishing a reference catalog of (pan)genomes (ideally from *de novo* genomes recovered from said dataset) and genome-specific pathway markers; 2) profiling this reference database using FASTQ reads; 3) aggregating counts relative to each pathway within each (pan)genome; and 4) estimating the “completeness” of essential steps within each pathway for each (pan)genome (i.e., pathway coverage). These (pan)genome-resolved functional abundance and coverage artifacts are the main outputs used for downstream analysis in ecosystem-level modeling.

Existing tools such as *HUMAnN* (6) have been instrumental in establishing a reproducible framework for functional profiling in metagenomics and metatranscriptomics. Recently, *Meteor2* (7) introduced a catalog-based approach that integrates taxonomic, functional, and strain-level profiling using ecosystem-specific microbial gene catalogs, improving upon existing methods for communities well-represented by pre-compiled catalogs. However, as sequencing costs decline faster than Moore’s Law predictions, the volume of meta-omic sequencing data has grown substantially and current profiling methodologies, whether catalog-based or alignment-dependent, are not designed for the new paradigm of genome-resolved metagenomics with massive custom genome collections (*AllTheBacteria* (∼2.7M genomes, https://allthebacteria.org/)*, GlobDB r232* (n=346,233 dereplicated genomes) (8)*, GTDB r232* (n=199,923 dereplicated genomes) (9), *OceanDNA* (n=52,325 genomes) (10), *Soil MAGs* (n=40,039 genomes) (11)). Furthermore, with a shift towards sample-specific genome-resolved metagenomics (12–14) and its dependence upon Average Nucleotide Identity (ANI) based pangenome clustering methods (12, 15, 16), the need for scalable profiling methods that can natively handle both genome- and pangenome-level resolution are essential for leveraging the power of rapid profiling techniques. With multiple strains per organism, pangenome-based approaches consider the entire gene repertoire of closely related genomes (e.g., strains clustered at a specific ANI and alignment fraction) and is essential for capturing the full functional landscape to understand functional redundancy or specialization within microbial populations (17).

Reference-based profiling in general can be performed on either a fixed reference catalog or a study-specific reference catalog. Precomputed fixed reference catalogs project every study onto the same feature axis, such as those used in the standard workflow of *HUMAnN* and *Meteor2*, so profiles can be compared across cohorts and environments. A study-specific reference, assembled *de novo* from the same or similar source samples, instead maximizes sensitivity to the organisms actually present and resolves function at the level of individual (pan)genomes recovered from that community at the cost of a feature axis particular to that study. Neither design is inherently more legitimate as they address different objectives. The advantage of study-specific references is working with the exact organisms in the study instead of inference by proxy to related strains making tangential results directly comparable to profiles such as genome quality, secondary metabolites, and antimicrobial resistance.

Custom genome-resolved reference catalogs for functional profiling are not novel. *Woltka* (18) derives taxonomically stratified functional profiles from alignments against custom genome collections and *MIDAS* (19) builds species-level pangenome databases from custom genome sets for intra-species gene content and variant analysis. However, *Woltka* operates over pre-computed alignments and does not address the alignment step itself while *MIDAS* targets intra-species diversity rather than pathway-level functional profiling. Neither computes per-(pan)genome pathway abundance and graph-based pathway completeness from a user-supplied reference in a single workflow, leaving this as an unmet need for researchers performing *de novo* genome-resolved metagenomics.

To address these challenges, we developed *Leviathan,* a rapid, modular, and scalable software package for integrated taxonomic and functional profiling of metagenomes and metatranscriptomes designed specifically for large *de novo* (pan)genome-resolved databases. *Leviathan* streamlines the workflows for 1) building taxonomic and functional profiling databases; 2) profiling taxonomic/sequence abundance; 3) profiling pathway abundance/coverage; and 4) merging sample-specific outputs lazily into *Xarray NetCDF* and *Apache Parquet* artifacts that can be seamlessly sliced into tabular dataframes.

This work introduces the architecture and capabilities of *Leviathan* by benchmarking runtime and memory usage against various backends and evaluating classification performance-using the standardized *CAMI I* (*Critical Assessment of Metagenome Interpretation*) low, medium, and high complexity datasets (20) and the marine *CAMI II* dataset (21). Finally, we demonstrate real-world case studies to stress-test *Leviathan* by analyzing a marine plastisphere metagenomics dataset and dental caries oral microbiome multi-omics dataset using metagenome-assembled genomes (MAG) with their source sequencing reads. These synthetic and real-world case studies demonstrate common paradigms in metagenomics while showcasing *Leviathan’s* scalability and native support for streamlined (pan)genome-level functional profiling representing a key advancement for comprehensive modern meta-omics research. *Leviathan* is an open-source software package available at https://github.com/jolespin/leviathan.

## Methods

### Benchmarking DNA-space and protein-space alignment performance on synthetic metagenomes

Benchmarking and analysis was performed on osx-arm64 using 4 threads running 3 jobs simultaneously for *Leviathan*, *Salmon*, *Bowtie2*, and *Diamond*. The following versions were used for all benchmarking: *Sylph v0.9.0*, *Salmon v2.3.1*, *Diamond v2.2.2, Leviathan v2026.7.16, VEBA v2.5.1,* and *PyKOfamSearch v2025.9.5*.

Computational performance benchmarks were determined by running *Salmon* backend using *Leviathan* configuration, *Bowtie2* backends using *HUMAnN* and *Meteor2* configurations, and *Diamond blastx* using *HUMAnN* configuration against the CAMI I and CAMI II synthetic metagenome datasets. Computational performance did not include *post hoc* filtering as the experiment was designed to unambiguously compare memory and runtime of various read assignment backends. Reference index and database build times were not considered during per-sample benchmarking.

To ensure fair comparison, each aligner was run using its documented or default paired-end read handling. *Salmon* received forward and reverse reads directly as used by *Leviathan*. For *HUMAnN’s Bowtie2* mode, paired-end reads were concatenated into a single FASTQ file and supplied as unpaired input (*-U*), following *HUMAnN* documentation. For *Meteor2’s Bowtie2* mode, forward and reverse reads were passed together as unpaired input *(-U R1,R2*), matching *Meteor2’s* prescribed usage. For *HUMAnN’s Diamond* mode, the concatenated FASTQ was converted to FASTA and used as input to *blastx*. For specific details on commands used, please refer to Supplemental Text S1.

Source genome- and pangenome-level alignment classification metrics were computed from mapped SAM and blast6 files with the following filtering strategies to reflect parent tool functionality: 1) *Leviathan*-configured *Salmon* alignments filtered by primary alignment only; 2) *HUMAnN*-configured *Bowtie2* alignments filtered by primary alignment and 50% reference gene coverage; 3) *Meteor2*-configured *Bowtie2* alignments filtered by primary alignment and ≥95% alignment identity; and 4) *HUMAnN*-configured *Diamond blastx* alignments filtered by ≥ 50% amino acid identity, e-value ≤ 1.0, and ≥ 90% query coverage. Since each backend resolves ambiguous mappings through fundamentally different mechanisms - expectation maximization (*Salmon*), pipeline-specific *post hoc* filtering (*HUMAnN*, *Meteor2*), or top-hit thresholding followed by *post hoc* filtering (*Diamond*) - primary alignments were used as the common evaluation unit for classification performance. This ensures that benchmarked performance reflects the alignment algorithm itself rather than downstream disambiguation strategies.

*Meteor2* involves post-filtering retaining reads ≥ 95% identity, loading all passing alignments into memory via dictionary keyed by read name to perform sort-order-agnostic best-hit selection. *HUMAnN’s Diamond blastx* backend uses FASTA-reformatted reads followed by multi-step filtering with proportional score redistribution across multi-mapped hits weighted by alignment quality. Cross-tool pathway-level benchmarking was not performed because the results would be confounded by annotation schemes, pathway definitions, and backend databases which cannot be directly compared.

In the case of this study, each sample-specific strain is considered a genome and all genomes from a dataset (e.g., *CAMI_medium*) are used to build pangenomes (e.g., genomes from *M2_S001* and *M2_S002* samples). For the purpose of this study, we define pangenomes as clusters of genomes with ANI ≥ 95% and minimum alignment fraction of 50% as used by *GTDB* (9, 34) and were computed using *VEBA’s* clustering module (12, 13). These genome and pangenome collections are used for building taxonomic and functional profiles. Co-assembly or pooled samples from the *CAMI* dataset were excluded to provide highly informative benchmarks with strain-level resolution. The set of CDS sequences with *KEGG* orthologs from pathways were used for building *Leviathan*, *Salmon*, and *Bowtie2* databases. The translated proteins from *Pyrodigal* for the same CDS set was used for building the *Diamond* database required for *Diamond blastx* benchmarking.

### Cross-catalog benchmarking against *OceanDNA*

To evaluate *Leviathan’s* behavior when the reference catalog is an independent, pre-compiled collection rather than study-specific MAGs, we profiled *CAMI-II* Marine reads against species-level representatives from the *OceanDNA* genome catalog (10). *OceanDNA* genomes were clustered at ANI ≥ 95% and alignment fraction ≥ 50% using *skani* (16), with the genome exhibiting the highest ANI connectivity within each cluster selected as the representative (12). CDS sequences from *OceanDNA* representatives were predicted with *Pyrodigal*, annotated with *PyKOfamSearch*, and used to build a *Salmon* index following the standard *Leviathan* workflow. *CAMI-II* Marine reads were then pseudo-aligned against this *OceanDNA* CDS catalog using *Salmon* with *Leviathan’s* default configuration.

Gold-standard genome assignments were established by computing pairwise ANI between all *CAMI-II* Marine source genomes and *OceanDNA* representatives using *skani*. A *CAMI* genome was considered “represented” if it matched an *OceanDNA* representative at ANI ≥ 95% and alignment fraction ≥ 50%, with one-to-one assignments enforced. Three metrics were computed: (1) catalog coverage, the fraction of *CAMI* source genomes with an *OceanDNA* representative; (2) accuracy of assigned reads, the fraction of mapped reads from represented genomes assigned to the correct *OceanDNA* target; and (3) false-positive rate, the fraction of reads from unrepresented genomes that received any assignment.

### Building the *Leviathan* reference index

The construction of a comprehensive reference index is a prerequisite for analysis with the *Leviathan* framework and is performed using the *`leviathan-index.py`* command-line executable module. This module requires a set of coding DNA sequences (CDS) in a FASTA file (*`--fasta`*), and a corresponding tab-separated feature mapping file (*`--feature_mapping`*) that links each gene ID to its functional annotations (as a Python-formatted set or list string) and its source genome ID. For convenience, *Leviathan* provides *`leviathan-preprocess.py`*, which takes as input a manifest file with (pan)genome identifiers, paths to genome assemblies, paths to CDS sequences, and annotations to reformat the data assets for *`leviathan-index.py`*. In addition, *`compile-manifest-from-veba.py`* is provided as a convenience to build the manifest files directly from *VEBA* outputs.

The module first validates data integrity by ensuring a perfect one-to-one correspondence between gene identifiers in the FASTA and feature mapping files before compiling the metadata into efficient, serialized Python dictionaries. For gene abundance/expression quantification, the module then builds a CDS-level index using *Salmon* (23), employing the *`--keepDuplicates`* flag to retain all provided sequences. This ensures that identical coding sequences carried by multiple genomes (e.g., from the same pangenome) are retained as distinct targets rather than collapsed, preserving genome provenance. *Salmon’s* expectation-maximization then apportions reads among shared targets such that the fractions sum to one. When all targets belong to the same pangenome, summing recovers exactly one count per read and when an equivalence class spans pangenome boundaries, the count is split across them.

Concurrently, if a mapping file of genome IDs to their assembly file paths is provided (*`--genomes`*), the module constructs a taxonomic profiling index by generating a single sketch of all genome assemblies using *Sylph* (1), with default parameters of a 31-bp k-mer, a minimum spacing of 30, and a subsampling rate of 200. If pangenome IDs are also provided, then the pangenome information is incorporated into the index and is used for downstream profiling. Pathway-level context is integrated by either using a pre-formatted custom database or by automatically downloading and processing the *KEGG* database if the provided functional features are identified as *KEGG* orthologs (24). The module ensures relevance by verifying that features in the pathway database overlap with those in the user’s dataset. The final output is a structured directory containing the *Salmon* index, the *Sylph* sketch, the compiled databases, configuration and log files, and a summary of MD5 hashes for all components. An existing *Leviathan* reference index can be updated *post hoc* to 1) include genome sketches by re-running the module with the *`--update_with_genomes`* flag or 2) update the *Salmon* backend database using the *`--update_salmon_index`* flag (e.g., migrating *Salmon* v1.x C++ database → *Salmon* v2.x Rust database). A typical command to build a new index is: *`leviathan-index.py --fasta pathway_markers.fasta[.gz] --feature_mapping gene_to_ko.tsv[.gz] --genomes genome_paths.tsv[.gz] --index_directory leviathan_index/ --n_jobs 8`*

### Taxonomic and sequence abundance with *Leviathan*

The taxonomic composition of a metagenomic/metatranscriptomic sample is determined using the *`leviathan-profile-taxonomy.py`* command-line executable module, which leverages the alignment-free *k*-mer-based profiler *Sylph* (1). The workflow begins by generating a *k*-mer sketch from the input paired-end sequencing reads (FASTQ format). This is handled internally by executing the *`sylph sketch`* command with default parameters of a 31-bp k-mer size (*`-k 31`*), a minimum spacing of 30 between selected *k*-mers, and a subsampling rate of 200 (*`-c 200`*); alternatively, a pre-computed reads sketch can be provided directly. The resulting reads sketch is then compared against the reference genome database (*`genomes.syldb`*) previously built and stored within the *Leviathan* reference index. The core profiling is performed by the *`sylph profile`* command, which estimates the relative abundance of each reference genome in the sample. To ensure high-confidence assignments, we used a minimum ANI threshold of 95% (*`--minimum-ani 95`*) requiring a minimum of 50 shared k-mers to consider a genome present (*`--min-number-kmers 50`*). The module also invokes the *`--estimate-unknown`* flag to account for the fraction of the community not represented in the reference database which is reported under the “`Sequence_abundance”` variable of the resulting artifacts.

Following profiling, *Leviathan* parses the *Sylph* output, converting genome file paths back to the user-defined genome identifiers from the index. It generates two primary metrics: “Taxonomic_abundance”, representing the relative proportion of each genome in the community, and “Sequence_abundance”, the fraction of total reads derived from each genome. If genome clusters (i.e., pangenomes) were defined during index creation, abundances are automatically aggregated by summing genome-level abundances by their pangenome mapping.

The final abundance tables are saved for each sample in either compressed TSV or *Apache Parquet* format at both the genome and pangenome levels. A typical command for taxonomic profiling is as follows: `leviathan-profile-taxonomy.py −1 forward.fastq.gz −2 reverse.fastq.gz - n Sample_1 -d path/to/leviathan_index/ -o output_directory/`

### (Pan)genome-resolved graph-based pathway completion and abundance with *Leviathan*

The functional potential and transcriptional activity of microbial communities is profiled using the *`leviathan-profile-pathway.py`* executable module, which integrates abundance/expression quantification with the genomic and pathway context stored in the *Leviathan* reference index. The workflow initiates by quantifying gene abundance/expression from paired-end sequencing reads.

Read assignment is performed using *Salmon* (23) in its `--meta` mode, which is specifically optimized for accurately assigning reads in complex metagenomic samples with strain-level heterogeneity similar to metazoan single-cell isoforms. To maintain high-confidence mappings, a stringent minimum alignment score fraction is used by default, ensuring that only reads with high-quality alignments contribute to abundance estimates. We swept *Salmon’s --minScoreFraction* parameter and found that micro-averaged F1 was stable across the tested range (0.65–0.95) while the fraction of gold-standard reads mapped decreased with stricter thresholds. We selected 0.87 as the default because it retains >95% of mapped reads relative to *Salmon’s* default (0.65) with no classification penalty; thresholds above 0.9 yielded no measurable improvement in micro F1 at substantial mapping cost, particularly for divergent communities (Figure S1, Table S1). However, *Leviathan* exposes this parameter so users can specify higher thresholds but *Leviathan*’s default follows that of *Alevin* for single-cell analysis which is designed to handle cell-type heterogeneity (25). The output of this step is a per-gene quantification table providing both raw read counts and Transcripts Per Million (TPM). As *Salmon* quantifies against the complete CDS catalog independently of *Sylph*, the functional module can assign reads to genes from genomes that the taxonomic module does not detect due to k-mer sampling, containment, or other parameter thresholds. Functional profiles can be filtered by taxonomic detection via indexing on the shared (pan)genome identifiers in the output *Parquet* or *Xarray* artifacts.

Following quantification, these gene-level abundances are contextualized by mapping each gene to its source genome and its annotated functional features (e.g., *KEGG* orthologs) using the metadata from the *Leviathan* reference index. The abundance/expression value of a gene annotated with multiple features is, by default, distributed equally among them, providing a scaled abundance for each feature but this can be turned off using the *--no_split_feature_abundances* option. A core feature of this module is its dual analysis of both functional potential or realized transcriptional activity for each pathway, resolved at the (pan)genome level.

*Leviathan* can calculate pathway completion through a graph-based approach via *KEGG Pathway Profiler,* if *KEGG* orthologs are provided as features, where each metabolic pathway is treated as a network of functional nodes (e.g., *KEGG* orthologs). *KEGG Pathway Profiler* can accept custom pathway databases but requires users to follow the same design schema. For each (pan)genome, its complete set of detected functional features are mapped onto these pathway graphs and a coverage score is computed representing the fraction of a pathway’s enzymatic steps encoded within that specific genomic context. *Leviathan* then measures the functional potential or transcriptional activity by aggregating the counts of all features constituting a given pathway within the same genomic unit. This yields a total pathway abundance value (in TPM or read counts) that reflects the collective abundance/expression of that pathway by a specific (pan)genome. When aggregating feature abundances to pathways, each feature contributes its full abundance to every pathway it participates in rather than being divided across them, as pathways represent an overlapping cover of feature space. This aggregation analysis is performed first at the individual genome level and if the *Leviathan* reference index was constructed with genome cluster definitions (i.e., pangenomes), then the module performs a subsequent aggregation, summing abundances and recalculating prevalence metrics for each cluster. More specifically, providing genome clusters as pangenome assignments covers the union span of the child elements including gene/feature-level and genome abundances as well as features that contribute to pathway coverage.

The final outputs are a series of structured tables (in Parquet or compressed TSV format) detailing gene, feature, and pathway metrics, including the dual measures of pathway abundance (activity) and coverage (potential), resolved at both the genome and, if applicable, the pangenome level. A typical command for the pathway profiling is as follows: `leviathan-profile-pathway.py −1 forward.fq.gz −2 reverse.fq.gz -n Sample_1 -d path/to/leviathan_index/ -o output_directory/`

### Merging sample-specific profiling into unified *Xarray NetCDF*

To facilitate study-level comparative analyses and downstream statistical modeling, *Leviathan* provides a dedicated *`leviathan-merge.py`* executable module that consolidates the per-sample outputs from the profiling workflows into unified, analysis-ready data structures. The inputs for this module are the parent directories containing the individual sample outputs from both the taxonomic and pathway profiling modules. It systematically discovers all per-sample result files (e.g., `.parquet` tables) and parses them to build comprehensive, multi-sample *N*-dimensional data structures that can be accessed via lazy loading without loading the entire file into memory.

The core of this process relies on the *Xarray* library (26), which is used to construct labeled, multi-dimensional arrays that preserve the relationships between samples, genomic units (genomes or pangenomes), and features (e.g., pathways, *KEGG* orthologs). This approach intelligently groups related metrics into single, coherent data objects. For instance, when merging pathway profiling results, the module generates a single *`pathways.genomes.nc`* file. This artifact contains an *`xarray.Dataset`* where distinct data variables for *`number_of_reads`, `tpm`,* and *`coverage`* share common dimensions for *`samples`, `genomes`,* and *`pathways`*. This structure is vastly more efficient for storage and analysis than managing separate flat files for each metric which is required for the scalability and performance needed for modern datasets.

A similar aggregation is performed for all other data types, including taxonomic abundances, feature abundances, and feature prevalence, at both the genome and pangenome levels. The final outputs are persisted as compressed *NetCDF* (`.nc`) files, a widely-used and highly scalable format. This data model simplifies complex data wrangling, prevents data misalignment, and provides a memory-efficient foundation for streamlined access to the entire project’s results, enabling robust, integrated multi-omic analyses.

### Pathway marker detection and completeness estimation

The standard workflow for *Leviathan’s* functional profiling engine uses: (i) *PyKOfamSearch* (https://github.com/jolespin/pykofamsearch) or *PyHMMSearch* (https://github.com/jolespin/pyhmmsearch), optimized implementations of *KofamScan* (24) and *HMMSEARCH* (27) utilizing *PyHMMER* (28) for rapid marker annotation; (ii) *Salmon* (23) for highly accurate and efficient pseudo-alignment of reads to gene catalogs for quantification; and (iii) the *KEGG Pathway Profiler* (https://github.com/jolespin/kegg_pathway_profiler) for robust, reaction graph-based assessment of KEGG pathway coverage and abundance. *KEGG Pathway Profiler* is an open-source reimplementation of *MGNify’s kegg-pathways-completeness-tool* (https://github.com/EBI-Metagenomics/kegg-pathways-completeness-tool) optimized for high-memory systems but could not have been developed without the fundamental work pioneered by EBI’s developers (29). *PyKOfamSearch*, *PyHMMSearch*, and *KEGG Pathway Profiler* have been developed to optimize annotation and are available within the *VEBA* software ecosystem (12). *Leviathan* expects pre-computed annotations so the user is able to provide annotations from any appropriate methodology and the standard workflow is only provided as a recommendation. *KEGG* compatibility is optional and users are expected to comply with *KEGG* licensing.

### Marine Plastisphere Microbiome differential abundance and coverage

Prokaryotic and eukaryotic genomes, pangenome-assignments, and protein-coding sequences from the *Marine Plastisphere Microbiome* case study were acquired from the official repository (FigShare: 10.6084/m9.figshare.20263974) and re-annotated with updated *KEGG* orthology via *PyKOfamSearch.* Genomes, pangenomes, and CDS sequences from annotated proteins were processed using *Leviathan*. *Leviathan’s* preprocessing, index, and pathway profiling modules were used to preprocess the metagenomes, generate a unified reference index, and profile pathways, respectively.

Differential abundance between early and late-stage plastic colonizing biofilms was performed at the pangenome-level with *leviathan profile-taxonomy* relative abundances via *ANCOM-BC* (*scikit-bio* v0.7.2) in categorical mode using a pseudocount of 1 to handle zeros during log-scale transformations (30, 31). The *ANCOM-BC* method accounts for varying sampling fractions between samples and addresses multiple testing with the Holm step-down method using Bonferroni adjustments for FDR correction of P-values (*statsmodels v0.14.6*, (32)). Statistical significance of differentially abundant pangenomes was determined using FDR < 0.001.

Differential coverage between early and late-stage plastic colonizing biofilms was performed at the pangenome-level with *leviathan profile-pathway* coverage values via two-sided Mann Whitney-U *test* (*scipy v1.17.1*) (33). Rank-biserial correlation was calculated from the Mann Whitney-U test statistic and used as a measure of effect size. Holm step-down method using Bonferroni adjustments was used to account for multiple testing when assessing differential abundance and differential coverage. FDR multiple test correction of P-values was implemented both at the pangenome-level and the more stringent global level across all features. Statistically significance of differentially covered pangenome-level *KEGG* modules was determined using FDR < 0.05.

### Dental Caries Oral Microbiome

Prokaryotic genomes, pangenome assignments, and protein-coding sequences from the *Dental Caries Oral Microbiome* case study were acquired from the official study repository (FigShare: 10.6084/m9.figshare.18180614.v1) and re-annotated with updated *KEGG* orthology via *PyKOfamSearch*. The genetic assets were processed through *Leviathan* following the methods described in the *Marine Plastisphere Microbiome* case study above. However, only the results of *leviathan profile-pathway* at the pangenome level were used for downstream analysis to demonstrate the utility of paired metagenomics and metatranscriptomics using *Leviathan*. Filtering of low-confidence features were filtered to only retain pangenome-level *KEGG* modules if they were 100% complete in at least 90% of the samples in either the caries-free or caries cohorts (*compositional v2023.8.28* (34)).

The *Differential Ensemble Co-expression Network* (DECN) was constructed using partial correlation with basis shrinkage as the co-expression metric (*n_draws* = 1000) with the caries-free cohort as the reference group and the caries cohort as the treatment group (35). Edges were pruned for both the positive (caries-enriched) and negative (caries-free enriched) connections using *BiDirectional Clustered Networks* (BDCN) which uses *Leiden* community detection, only retaining edges that cluster together for 100% of the 1000 iterations using separate random seeds (36). Both DECN and BDCN were computed via the *ensemble_networkx v2025.7.8* Python package (37).

Positive and negative networks with signed edge weights were merged for visualization and network property calculations such as connectivity (summed connection weight) and betweenness centrality (shortest-path betweenness centrality for nodes) calculated via *NetworkX v3.6.1.* Nodes with disproportionately high betweenness-centrality were determined by using a robust adaptation of z-score as a threshold: *Median* + 3 *Median Absolute Deviance* (MAD) via *scipy*.

## Results

### Overview of *Leviathan*

*Leviathan* is an open-source, command-line-driven software package designed for rapid, scalable, and integrated taxonomic and functional profiling of metagenomic and metatranscriptomic data. The entire workflow is engineered for high performance and centered around a modular, three-stage architecture that streamlines analysis from FASTQ reads to multi-sample, analysis-ready data artifacts (Figure 1). *Leviathan* is run at the sample level, optimized for distributed compute environments, and the individual tabular artifacts (.parquet or .tsv) are merged into *Xarray NetCDF* files for lazy loading and efficient access to large structured database tables.

**Figure 1.**
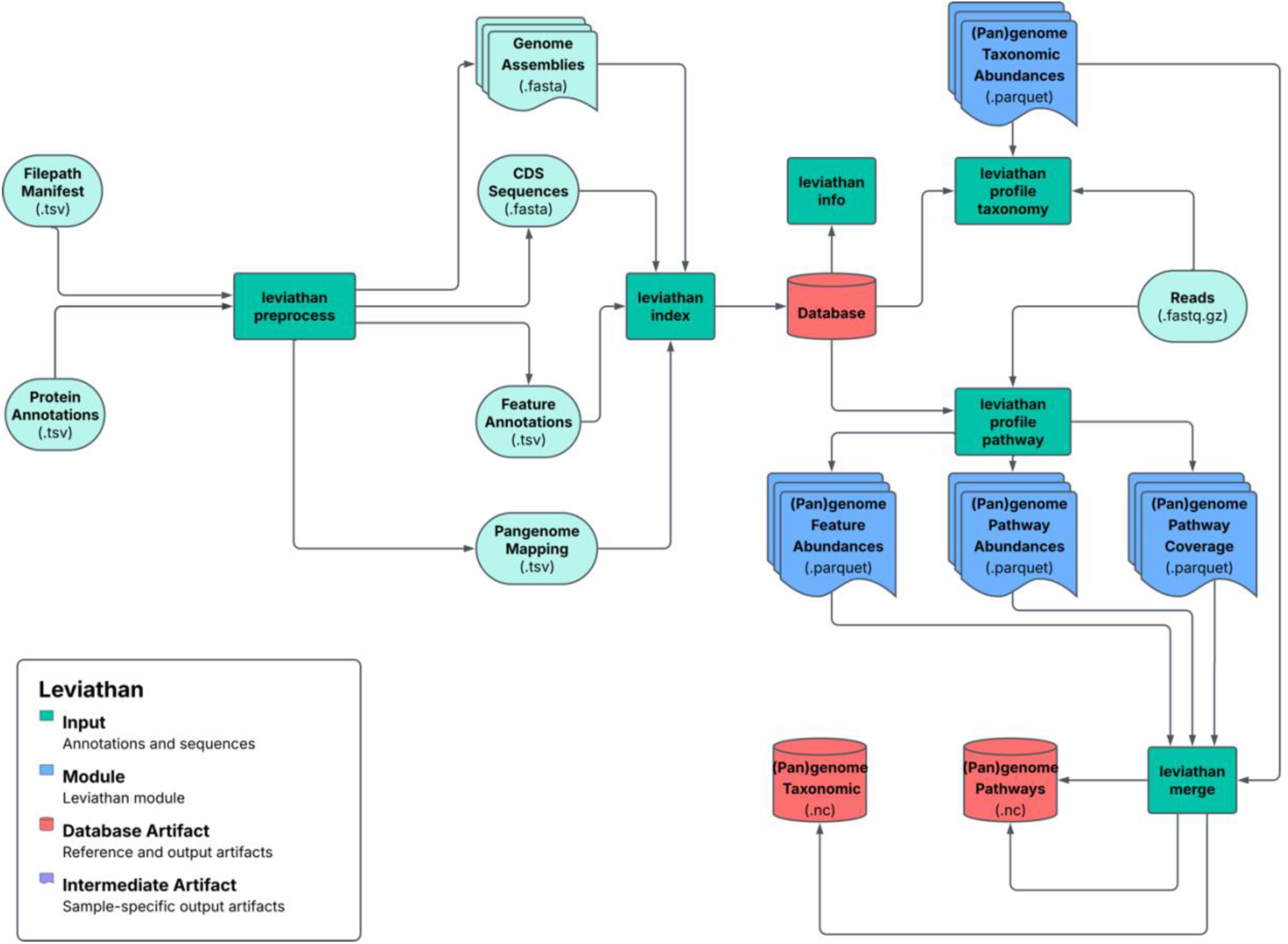
Flowchart of *Leviathan* core modules

The first stage is database construction via *`leviathan-index`*. *Leviathan* builds a single, unified reference index containing a *Sylph* sketch for *k*-mer-based taxonomic profiling, a *Salmon* index of coding sequences for high resolution functional quantification, and pre-compiled databases that map genes to functional features (e.g., *KEGG* orthologs) and pathways. Optionally, genome clusters (i.e., pangenomes) can be provided during the database construction which will be used automatically in profiling stages. This step efficiently prepares all necessary assets for downstream analysis. While we use ANI and alignment fraction-based pangenome clustering, *Leviathan* supports custom clustering approaches or dereplication methods (38)

The second stage is profiling with taxonomic or functional profiling which are independent of each other. Taxonomic profiling via *`leviathan-profile-taxonomy`* is essentially a convenient wrapper around *Sylph* to rapidly estimate the relative abundance of reference genomes in a sample. The core innovation of *Leviathan* exists in its functional profiling module via *`leviathan-profile-pathway`*, which first quantifies gene-level abundance/expression using *Salmon’s* meta-mode for high accuracy in complex communities. Unlike traditional aligners that compute base-level read-to-reference alignments, *Salmon’s* pseudo-alignment determines read origin by mapping k-mer compatibility classes to reference sequences, bypassing positional alignment entirely while using expectation-maximization to resolve ambiguous assignments. After pseudo-alignment, *Leviathan’s* functional module computes two distinct metrics for each pathway at the (pan)genome level: 1) pathway coverage, a graph-based assessment of the completeness of a pathway’s enzymatic steps, representing its genomic potential; and 2) pathway abundance, the aggregated abundance/expression of all genes within that pathway, representing its functional potential/activity. Both taxonomic and functional profiling backends (i.e., *Sylph* and *Salmon*) are implemented in Rust for ultra-fast performance. Although taxonomic and functional profiling are independent and can be run separately, a user can easily choose to filter functional profiles by taxonomic abundances or vice versa via indexing on downstream parquet or *Xarray* data structures.

The final stage is merging the artifacts into individual data structures via *`leviathan-merge`*. *Leviathan* consolidates the individual per-sample outputs into individual multi-dimensional *Xarray NetCDF* files. This creates a memory-efficient, lazy-loading supported analysis-ready data structure that contains all metrics (e.g., taxonomic abundance, sequence abundance, pathway coverage, pathway abundance (number of reads), pathway abundance (tpm)) conveniently organized by sample, feature, and genomic unit. A key design principle throughout this workflow is *Leviathan’s* native support for pangenome-resolved analysis, where all metrics can be seamlessly aggregated from individual genomes to user-defined pangenome clusters.

### Native support for genome and pangenome-level profiling of microbial communities

Next-generation sequencing instruments produce data that is inherently compositional as they estimate the relative abundance of discrete biological components (e.g., transcripts, marker genes) within a community by sampling from a pool of nucleic acid fragments (34). Biological features are naturally hierarchical (e.g., genes → chromosomes/plasmids → genomes → pangenomes) and individual components can be amalgamated up the hierarchy while retaining compositionality (40). In the case of *Leviathan*, the relative abundance and counts tables produced by *Sylph* and *Salmon* in the backend, respectively, are compositional and, thus, individual component abundances can be aggregated with respect to higher order biological structures such as pangenomes and pathways. In the context of this study, a strain will be referred to as a genome and each genome is recovered from an individual sample. Each output artifact contains a genome and pangenome version allowing users to easily select which level would be appropriate for their analysis. We refer to the pathway-level output features as Taxa-Resolved Functional Modules (TRFM) where each TRFM represents the coverage and abundance of a specific metabolic pathway within a single genome or strain-dereplicated pangenome.

One caveat for estimating the abundance of a feature or pathway is that a single gene can be assigned to multiple features (e.g., *KEGG* orthologs or enzymes) and a feature can be represented in multiple pathways. To address this, *Leviathan* implements a fractional abundance for these scenarios which can be toggled off. An example of this would be *Salmon* mapping a read to *gene_a* which is annotated by 3 features {*KOfam_1, KOfam_2, KOfam_3*}. When aggregating the gene-level counts with respect to each feature, the read count would be equally distributed across the features where each feature would receive ⅓ of a count for each read mapped to *gene_a* (or ⅓ of a TPM value if using TPM as a metric). This approach ensures that the sum of feature counts equals the sum of gene counts which is necessary for interpreting relative abundances for compositional data analysis.

### *Leviathan* integrates *Sylph* for rapid (pan)genome-resolved taxonomic profiling

*Leviathan* provides users with *leviathan-profile-taxonomy,* a convenient wrapper around *Sylph* which is an ultrafast and precise taxonomic profiler (1). *Sylph* is a species-level metagenome profiler that estimates genome-to-metagenome containment ANI through zero-inflated Poisson *k*-mer statistics and has superior performance relative to other approaches with support for genomes at low coverage levels (See Shaw and Yu, 2024 for benchmarking details). After running *Sylph*, *Leviathan* sums the taxonomic and sequence abundance values with respect to pangenomes and outputs relative abundance tables for genomes and pangenomes. Once the taxonomic profiling has been completed for each sample, the abundance tables are merged to build *Xarray NetCDF* artifacts where users can access data using lazy loading without storing everything into memory.

### DNA-space and protein-space alignment performance on synthetic metagenomes

*CAMI* datasets are synthetic metagenomics datasets widely adopted for gold-standard benchmarking of metagenomic tools (18, 19). For benchmarking, we have included the *CAMI-I low complexity* (S:1, G:30, P:30), *CAMI-I medium complexity* (S:2, G:450, P:270), *CAMI-I high complexity* (S:5, G:2036, P:789), and *CAMI-II marine* (S:10, G:6359, P:2647) where S, G, and P refer to the number of unique samples, genome assemblies, and pangenomes in each dataset. *CAMI* provides ground-truth tables for read → contig → genome → sample) allowing for accuracy-based benchmarking of read-alignment backends.

Comparing profiling tools end-to-end conflates multiple variables: database composition, annotation scheme (e.g., *UniRef* (41) with transitive *KEGG* mapping vs. *KOfam* directly), and post-alignment processing. To isolate the read-alignment step, the primary computational bottleneck and the locus of the DNA-space vs. protein-space tradeoff, we benchmarked four backends against a common CDS catalog derived from each *CAMI* dataset’s reference genomes: (1) *Salmon* pseudo-alignment using *Leviathan’s* configuration, (2) *Bowtie2* alignment using *HUMAnN’s* configuration, (3) *Bowtie2* alignment using *Meteor2’s* configuration, and (4) *Diamond blastx* translated search using *HUMAnN’s* configuration. Classification performance was evaluated at both genome and pangenome levels. Within a single dataset there are multiple samples and each sample may or may not contain a separate strain of an organism.

*CAMI I* and *II* datasets include 8875 genomes that cluster into 3736 pangenomes (Table 2). Of the 8875 genomes, 856 genomes were insufficient for *Pyrodigal* to identify any protein-coding genes, 3129 genomes were medium-to-high quality (*CheckM2* completeness ≥ 50% and contamination < 10%), and 7202 genomes contained robust *KEGG* pathway *KOfam* annotations. *Leviathan* functional profiling can only incorporate genomes with feature annotations (e.g., *KEGG* pathway *KOfam* in this study), therefore, any genome that did not have *KOfam* annotations in a *KEGG* pathway that exceeded the HMM threshold were excluded from the analysis. However, every genome that lacked these annotations also failed medium stringency quality control based on *CheckM2* so they would be excluded from best-practice workflows. A separate reference catalog for each dataset was built using the subset of genes (or translated proteins) with orthology to *KEGG* pathway *KOfams* and used across backends for direct comparisons.

**Table 1.** Comparison of features between *Leviathan,HUMAnN,* and *Meteor2*. N.D indicates that usage is not documented on the official repository.

| Feature | *Leviathan* | *HUMAnN* | *Meteor2* |
| --- | --- | --- | --- |
| Alignment method | *salmon quant* | *bowtie2*, *diamond*<br>*blastx* | *bowtie2* |
| Alignment space | DNA | DNA, Protein | DNA |
| Requires *MinPath* | No | Yes | No |
| Requires taxonomic lineage | No | Yes | No |
| Functional annotational method | User-provided (e.g., HMM with score thresholds) | Transitive (*Read* → *UniRef* → *MetaCyc*) | Catalog-derived |
| Supports custom genome catalogs | Yes | Yes, but custom pangenomes requires building *MetaPhlAn* + *ChocoPhlAn* databases | N.D. |
| Supports custom annotations | Yes | Yes (*UniRef*) | N.D. |
| Supports custom pathways | Yes | Yes | N.D. |
| Supports functional profiling | Yes | Yes | Yes |
| Supports pangenomes | Yes | Yes | Yes |
| Supports single-end reads | Yes | Yes | Yes |
| Supports paired-end reads | Yes | No, requires concatenation | Partially, treated as unpaired |
| Supports long reads | No | No | No |
| Supports taxonomic profiling | Yes | No | Yes |
| Supports stratified counts tables | No | Yes | No |
| Supports lazy loading data structures | Yes | No | No |

**Table 2.**
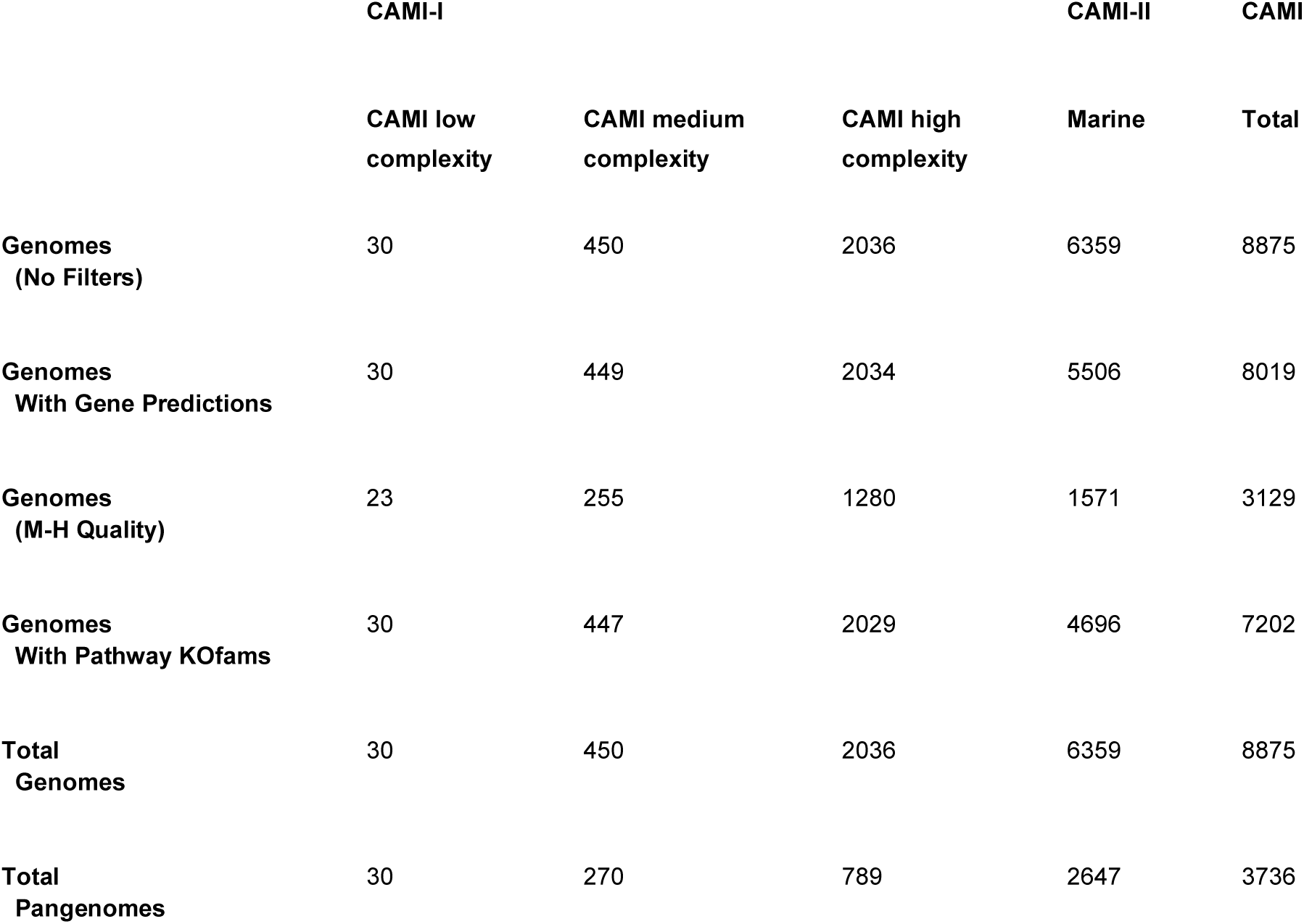
Comparison of genomes from CAMI datasets.

The three DNA-space backends *Salmon*, *Bowtie2-HUMAnN*, and *Bowtie2-Meteor2* completed read assignment with a median runtime of 2.45 minutes, 9 minutes, and 5.16 minutes using peak memory usage of 1.06 GB, 1.07 GB, and 1.13 GB, respectively while *Diamond blastx* (*HUMAnN* config) required a median of 91.92 minutes and 12.8 GB (Fig. 2A). *Salmon was* faster than both *Bowtie2*-*HUMAnN* configuration (--very-sensitive), *Bowtie2-Meteor2* configuration (--sensitive) as well as the *Diamond blastx HUMAnN* configuration by a median fold improvement of 3.42x, 2.14x, and 34.58x. respectively. *Salmon* used similar memory to both *Bowtie2* configurations and DNA-space aligners used considerably less than *Diamond blastx*. Fold improvements were computed within-sample and then summarized because runtime spans an order of magnitude across datasets, ratios of pooled medians would be confounded by dataset complexity.

**Figure 2.**
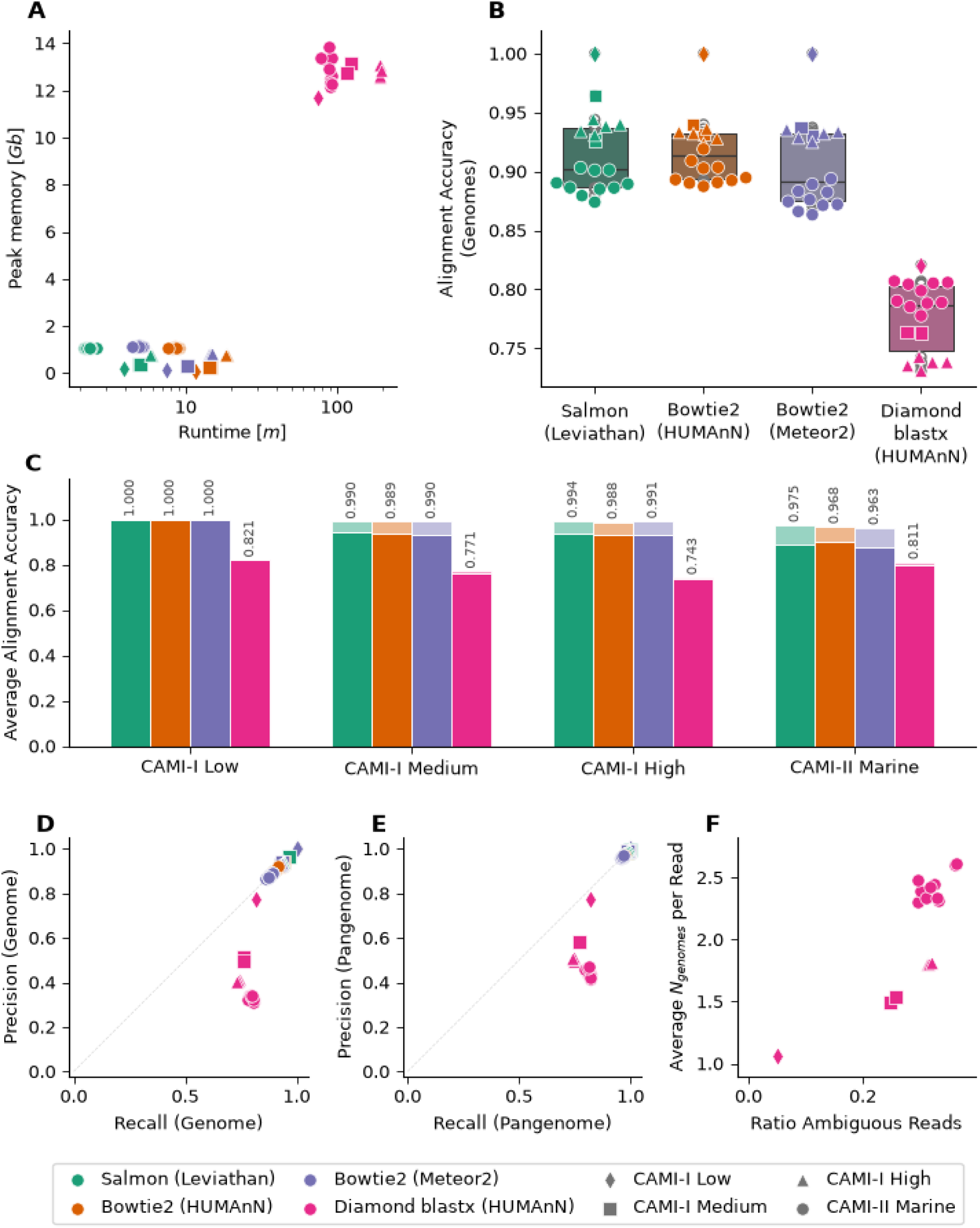
Read-alignment backend benchmarking against *CAMI* gold-standard datasets. Four backends were evaluated: *Salmon* (DNA-space pseudo-alignment, *Leviathan* configuration), *Bowtie2* using *HUMAnN* configuration (nucleotide alignment), *Bowtie2* using *Meteor2* configuration/post-filtering (DNA-space alignment), and *Diamond blastx* using HUMAnN configuration/post-filtering (translated protein-space search). Marker shape denotes the dataset where each point represents an individual sample. Computation performance showing (A) peak memory in gigabytes versus runtime in minutes (log scale) for each backend on each dataset using 4 threads each. (B) Distribution of genome-level alignment micro-averaged accuracy across samples grouped by backend. (C) Mean accuracy per dataset where each bar is partitioned into genome-level accuracy (solid) and the additional fraction of reads scored correct when the criterion is relaxed to pangenome-level assignment (opaque) annotated with pangenome-level accuracy. Means are used instead of medians for stacked bars as decomposition into genome-level accuracy plus the pangenome-level increment is additive only under the means. (D, E) Micro-averaged precision versus recall at the genome (D) and pangenome (E) levels; the dashed line marks precision = recall. (F) Protein-space multi-mapping behavior for *Diamond blastx*, showing mean number of genome assignments per read versus the fraction of reads mapped to more than one genome. *Diamond blastx --top 1* configuration retains multiple hits per read as *HUMAnN’s* downstream processing expects. DNA-space backends are omitted as primary alignment filtering yields exactly one assignment per read.

Alignment accuracy at the genome level, the fraction of reads assigned to the correct source genome, was comparable across all three DNA-space backends, with median micro-averaged accuracies of 90.2%, 91.4%, and 89.1% for *Salmon*, *Bowtie2-HUMAnN*, and *Bowtie2-Meteor2*, respectively (Fig. 2B). *HUMAnN’s* implementation of *Diamond blastx* uses *--top 1* argument which allows multiple hits per read where read counts are distributed across spread but these hits can map to different genomes (Fig. 2F). *Diamond blastx* achieved a lower median accuracy of 78.7% using relaxed scoring (i.e., any correct hit), 62.4% accuracy using weighted scoring (i.e., 1/n where n is number of unique genome hits per read), and 54.9% accuracy using stringent scoring (i.e., all correct hits) with the greatest disparity observed in the *CAMI-I* high complexity dataset (Fig. 2C, Table S1). This accuracy reduction is consistent with the behavior of translated search against genome-resolved CDS catalogs: protein-space alignment introduces additional ambiguity when closely related genomes share conserved protein sequences that are distinguishable at the nucleotide level.

The residual precision gap for *Diamond blastx* after pangenome aggregation (34.2% → 47.1%) is explained by multi-mapping behavior intrinsic to protein-space search (Fig. 2F). *Diamond blastx* --top 1 configuration yields 5% ambiguous reads with a mean of 1.06 genome assignments per read in *CAMI-I* low complexity and 32% ambiguous reads with a mean of 2.4 genome assignments per read in *CAMI-II* Marine (Fig. 2F, Table S1). In protein-space, synonymous nucleotide variants collapse into identical protein sequences, causing reads from conserved coding regions to match proteins from multiple source genomes across pangenome boundaries, introducing a source of ambiguity that pangenome-level aggregation cannot resolve. DNA-space backends are omitted from this panel as pangenome-level aggregation recovered their precision to ≥96.8% (Fig. 2E), leaving no residual gap to diagnose; each read receives a single primary assignment and the remaining genome-level error is attributable to within-species strain ambiguity resolved at the pangenome level.

Classification performance was further evaluated using precision and recall at both genome and pangenome levels (Fig. 2D,E). For the DNA-space backends, each read receives a single genome assignment and under micro-averaging every misassignment is simultaneously a false positive for one genome and a false negative for another, yielding identical precision and recall values. At the genome level, DNA-space backends achieved micro-averaged precision and recall of 90.2% (*Salmon*), 91.4% (*Bowtie2-HUMAnN*), and 89.1% (*Bowtie2-Meteor2*). *Diamond blastx* breaks this symmetry because the --top 1 parameter permits multiple hits per read across different genomes. Each additional incorrect genome hit contributes a false positive without a corresponding false negative, inflating the denominator of precision while recall remains relatively high as the correct genome is typically retained among the hits. This produced a median genome-level precision of 34.2% against a median recall of 78.6% for *Diamond blastx* (Fig. 2D).

At the pangenome level, aggregation within species-level genome clusters resolved the majority of classification errors across all DNA-Space backends (Fig. 2C-E) with near-perfect accuracy (Fig. 2C). DNA-space backends achieved pangenome precision and recall median values of 98.1% (*Salmon*), 97.3% (*Bowtie2-HUMAnN*), and 96.8% *(Bowtie2-Meteor2*), demonstrating that genome-level misassignments are predominantly between strains of the same species rather than between unrelated taxa. *Diamond blastx* pangenome precision improved from 34.2% to 47.1%, indicating that within-species multi-mapping is partially resolved by aggregation but a residual gap persists from multi-mapped reads spanning pangenome boundaries (Fig. 2F).

Across all CAMI datasets, *Leviathan* completed end-to-end functional profiling in a median of 5.26 minutes per sample with a peak memory usage of 2.8 GB (Table S1). For context, this total pipeline cost includes pseudo-alignment, feature mapping, fractional abundance estimation, and graph-based pathway coverage computation. The classification metrics for *Leviathan’s* functional profiling are identical to *Salmon* benchmarks as this is used as the backend. At this per-sample cost, functional profiling of large cohort studies can be trivially parallelized across samples without specialized hardware requirements.

To assess whether *Leviathan’s* alignment score threshold effectively prevents spurious assignments when profiling against a reference catalog that is independent of the study, we benchmarked *CAMI-II* Marine reads against *OceanDNA* species-level representatives. Of the 6,359 *CAMI-II* Marine source genomes, 403 (6.3%) had an *OceanDNA* representative at ≥ 95% ANI (Fig. S2, Table S1). Among reads from represented genomes that received a *Salmon* assignment, 94.2% were assigned to the correct *OceanDNA* target genome, with a per-genome median accuracy of 97.7%. Reads from the 5,956 unrepresented genomes, that is, *CAMI-II* Marine genomes with no species-level match in *OceanDNA*, had false-positive assignment rate of 0.2%, with a median per-genome rate of 0.0% (Fig. S2). These results demonstrate that *Leviathan’s* default alignment score threshold prevents spurious assignments to non-target genomes while maintaining high accuracy for organisms with catalog representation.

### Case Study I: Marine Plastisphere Microbiome

The *Marine Plastisphere Microbiome* metagenomics cross-sectional dataset represents microbial communities from early and mature stage biofilms on plastics in marine surface water (13, 42). Of these samples, 17 early stage biofilms were collected from virgin plastic particles incubated in seawater for 2-7 days while 20 late stage biofilms were sourced from plastic litter harvested at the same locations. Genome-resolved metagenomic analysis yielded 219 prokaryotic and 5 eukaryotic MAGs clustering into 154 prokaryotic and 4 eukaryotic pangenomes. For more details regarding the experimental setup, see Bos et al., 2023.

To showcase the utility of *Leviathan* for metagenomics, we first investigated which microbes were enriched separately in early and mature stage plastic forming colonies. Differential abundance tests revealed 19 taxa and 6 taxa enriched in early-stage and mature-stage plastic biofilms, respectively (Fig. 3, Table S2). Taxa from the family *Alteromonadaceae* were highly enriched in early-stage plastic colonies compared to mature-stage plastic colonies. Of the top 6 highly enriched taxa in the early-stage colonization, 4 were of the *Alteromonadaceae* family, *Marisediminitalea aggregata* (log_2_FC = −8.14) and *Alteromonas macleodii* (log_2_FC = −6.60) exhibiting the greatest bias-corrected log-transformed abundance, while the other enriched taxa included *Marinobacter flavimaris* (log_2_FC = −4.5) and an uncharacterized *Saccharospirillaceae* (log_2_FC = −4.3). Mature-stage plastic samples were dominated by *Alphaproteobacteria* including an uncharacterized *Hyphomicrobiaceae* (log_2_FC = 4.16), an uncharacterized *Rhodobacteraceae* (log_2_FC = 4.1), an uncharacterized *Ahrensia* (log_2_FC = 3.95), an uncharacterized *Erythrobacter* (log_2_FC = 3.34), and an uncharacterized *Henriciella* (log_2_FC = 3.02), in addition to an uncharacterized *Saprospiraceae* with slightly lower enrichment (log_2_FC = 2.44). The differential abundance results are consistent with the plastisphere succession patterns described by Bos et al. 2023, with Alteromonadaceae-dominated copiotrophic Gammaproteobacteria in early-stage biofilms giving way to Alphaproteobacteria-dominated mature biofilm communities, including Rhodobacteraceae and Hyphomonadaceae (the taxonomic family of *Henriciella)*.

**Figure 3.**
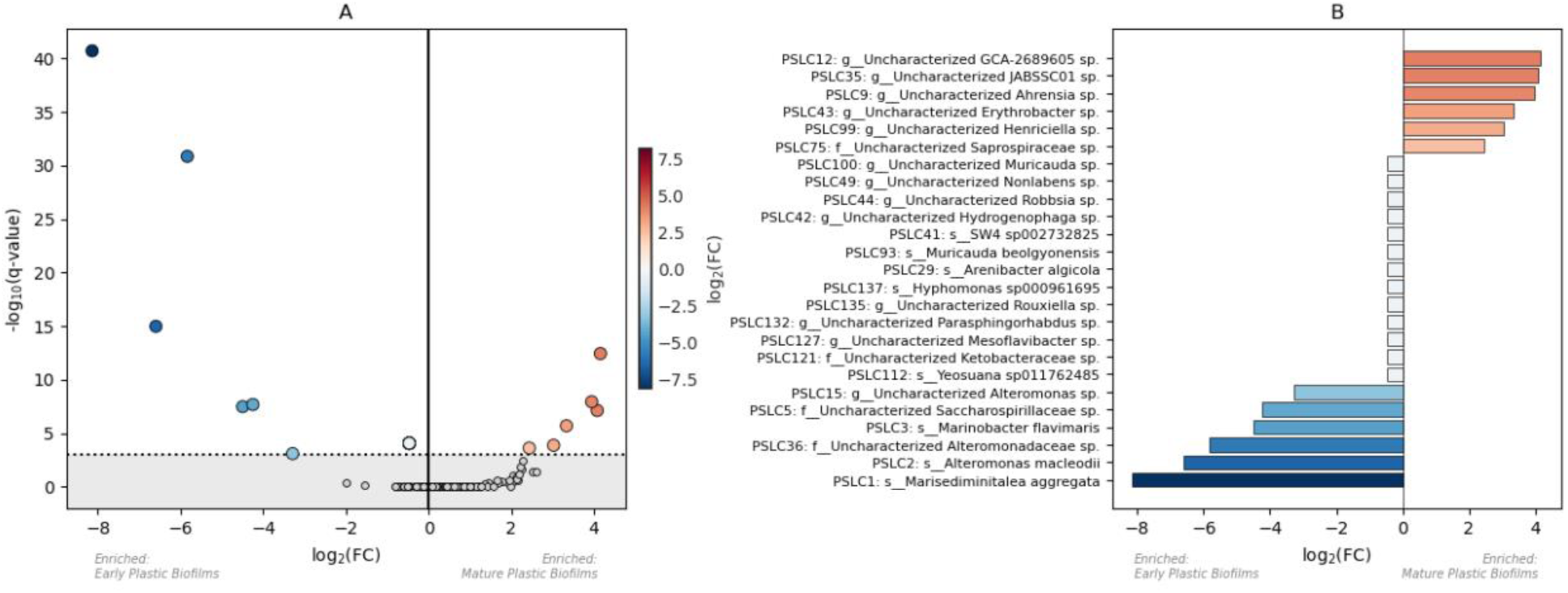
Marine plastisphere microbiome case study results from *ANCOM-BC* differential abundance test of pangenome-level taxonomic abundance. (A) Volcano plot with data points representing pangenomes (N=158) and dotted line denoting the global significance threshold (FDR < 0.001). Data points above significance threshold are enriched in the early (left) or mature (right) plastic biofilms with (B) ANCOM-BC Log2FC values shown in barchart. Taxonomic labels represent the lowest resolved rank from the consensus taxonomy of each pangenome.

To assess the variation of metabolic potential of pangenomes between early and mature-stage plastic colonizing microbes, we leveraged the functional profiling results using TRFM coverage profiles from the *leviathan profile-pathway* module at the pangenome level as they represent the metabolic capability of strain-dereplicated organisms. *Leviathan’s* functional profiling for the plastisphere metagenomics dataset had a median runtime of 1.65 minutes and peak memory of 1.85 GB (Table S1). While the abundance of specific functional modules from metagenomics would roughly approximate the abundance of microbes, coverage can vary as microbes lose or gain genes through evolution or horizontal gene transfer. To complement the differential taxonomic abundance, we performed differential coverage analysis to investigate which TRFMs differed statistically in detected pathway step completion between early and mature plastic biofilm communities.

We identified 664 TRFM and 30 TRFM with statistically higher *KEGG* module completion ratios in early and mature-stage plastic colonizers, respectively (Fig. 4, Table S3). Early-stage taxa, primarily dominated by *Alteromonadaceae*, exhibited broadly higher *KEGG* module completion across cofactor/vitamin metabolism (Pimeloyl-ACP biosynthesis) and central carbohydrate metabolism (gluconeogenesis and glycolysis) while mature-stage taxa exhibited higher *KEGG* module completion across amino acid metabolism (serine/threonine, lysine, and branched-chain metabolism), central carbohydrate metabolism (citrate cycle and pyruvate oxidation), and cofactor/vitamin biosynthesis (Pantothenate and Coenzyme A biosynthesis). These differential coverage patterns complement the taxonomic abundance results, demonstrating how *Leviathan’s* pangenome-level functional profiling can reveal metabolic plasticity that would be obscured at the individual genome level.

**Figure 4.**
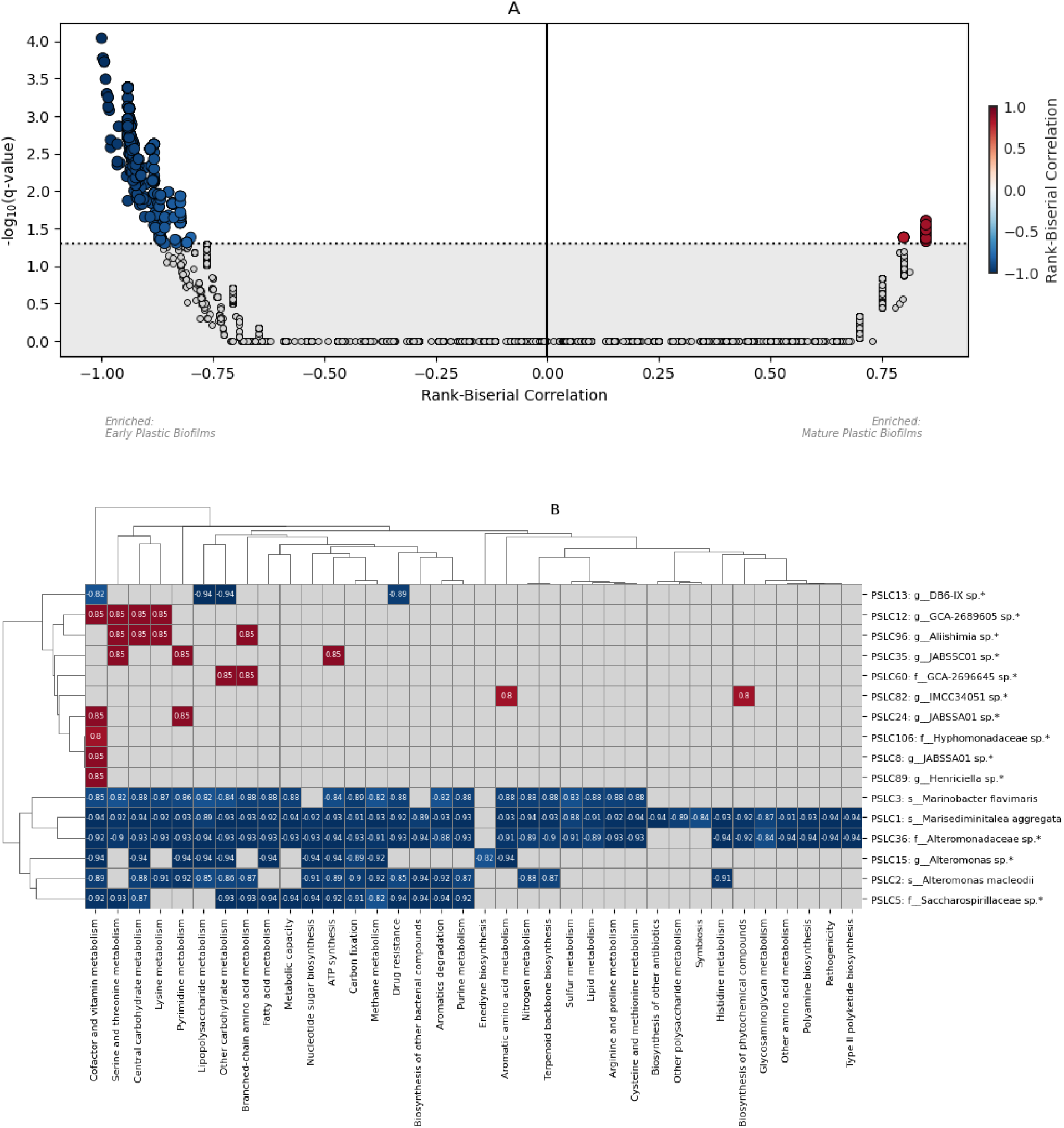
Marine plastisphere microbiome case study results from Mann Whitney-U differential pathway coverage test of pangenome-level with rank-biserial correlation. Positive (red) values and negative (blue) values indicate rank-biserial enrichment in mature-stage and early-stage biofilms, respectively. (A) Volcano plot with data points representing pangenome-specific *KEGG* modules (N=25,263) and dotted line denoting the global significance threshold (FDR < 0.05). Data points above significance threshold are enriched in the early (left) or mature (right) plastic biofilms with (B) rank-biserial correlation values averaged per pathway category shown in clustermap.

### Case Study II: Dental Caries Oral Microbiome

The *Dental Caries Oral Microbiome* multi-omics cross-sectional dataset represents microbial communities from supragingival plaque of Australian twin children. This study recovered 658 prokaryotic MAGs that clustered into 135 pangenomes. These genetic assets were used to build a *Leviathan* reference database and the associated 91 metatranscriptomes were leveraged to assess transcriptional patterns in caries and caries-free cohorts (35, 43). *Leviathan’s* functional profiling for the oral metatranscriptomics dataset had a median runtime of 1.31 minutes and peak memory of 1.37 GB (Table S1).

To showcase the utility of *Leviathan* for multi-omics (metagenomics and metatranscriptomics) from an ecological perspective, we built a compositionally-valid DECN using caries-free and caries as the reference and treatment cohorts, respectively, then clustered these networks into BDCNs. Nodes in this network represent high-confidence TRFM expression (i.e., pangenome-resolved *KEGG* modules that were 100% complete in at least 90% of the cohort) and edges represent coexpression of functional modules between organisms. While co-expression of TRFM is not a replacement for robust yet resource intensive community-scale metabolic models (44), co-expression can serve as a proxy for characterizing broad transcriptional patterns that are indicative of microbial communities in the context of health and disease.

The caries-enriched subnetwork clustered into 12 Leiden communities ranging from 2-12 nodes per cluster including a total of 53 nodes and 198 edges representing 24.5% and 1.71% of initial DECN nodes and edges, respectively. The caries-free-enriched subnetwork clustered into 18 Leiden communities ranging from 2-21 nodes per cluster including a total of 108 nodes and 489 edges representing 50% and 4.2% of initial DECN nodes and edges, respectively. The union of the caries and caries-free-enriched networks includes 142 nodes with 19 intersecting nodes indicating distinct metabolism within communities and a low proportion of bridge nodes relevant to both caries and caries-free transcriptomic states (Fig. 5, Tables S4,S5). The BDCN algorithm asserts that no edges overlap between the caries-enriched and caries-free-enriched clustered networks.

**Figure 5.**
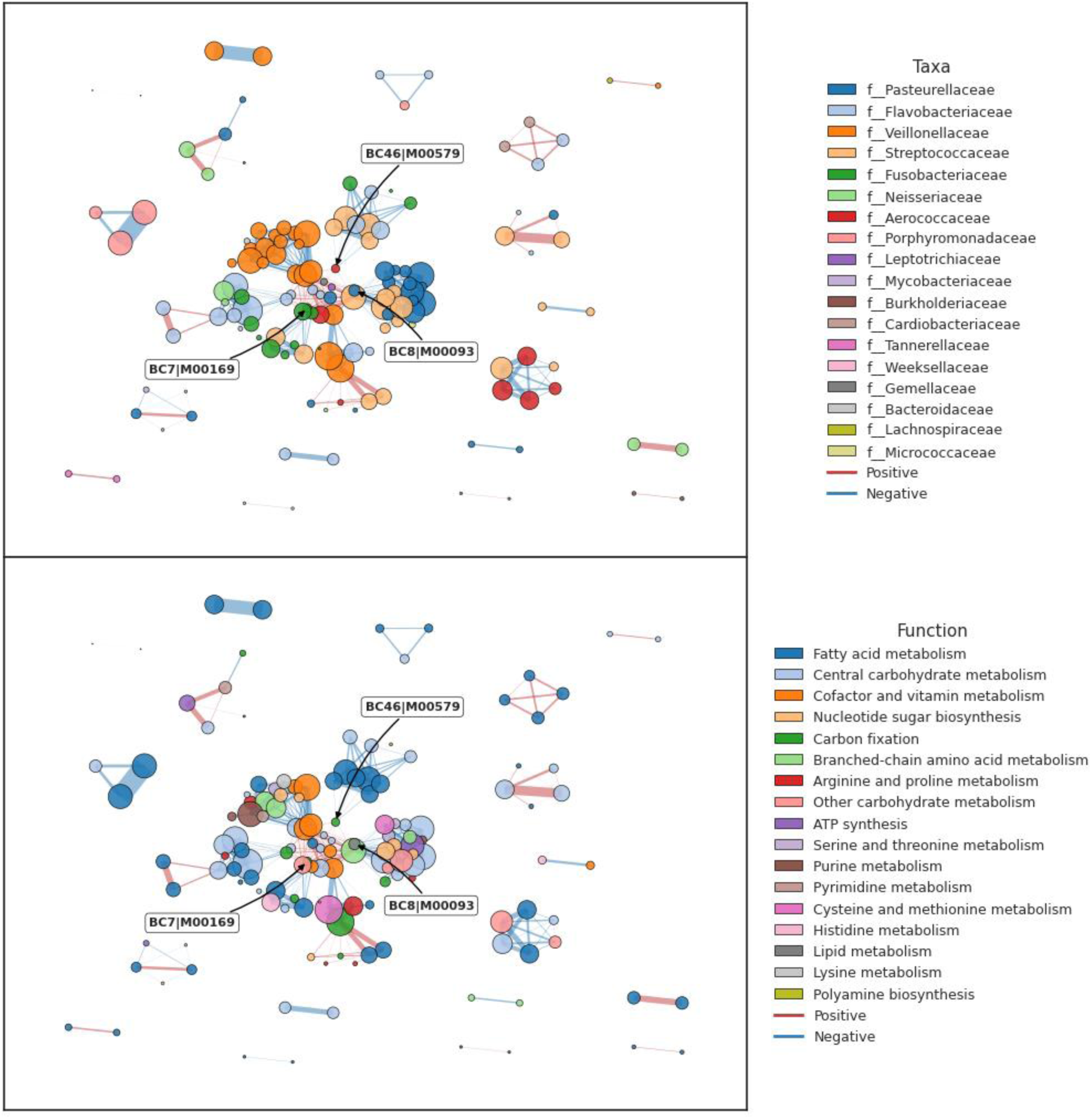
BiDirectional Clustered Networks TRFM connectivities enriched in caries cohort (red) and caries-free cohort (blue). TRFM nodes colored by (A) taxonomy and (B) pathway category.

The union BDCN reveals bridge nodes connecting modules dense with caries-enriched connections to modules dense with caries-free-enriched connections as shown in Fig. 5, S3 We used betweenness-centrality (BC) to assess which bridge nodes to interpret for demonstrating the utility of *leviathan profile-pathway* in the context of investigating caries from an ecological perspective (Table S5). Our robust adaptation of the standard z-score uses median and MAD instead of mean and standard deviation, making it resistant to outliers and therefore better suited for detecting them (Fig. 6).

**Figure 6.**
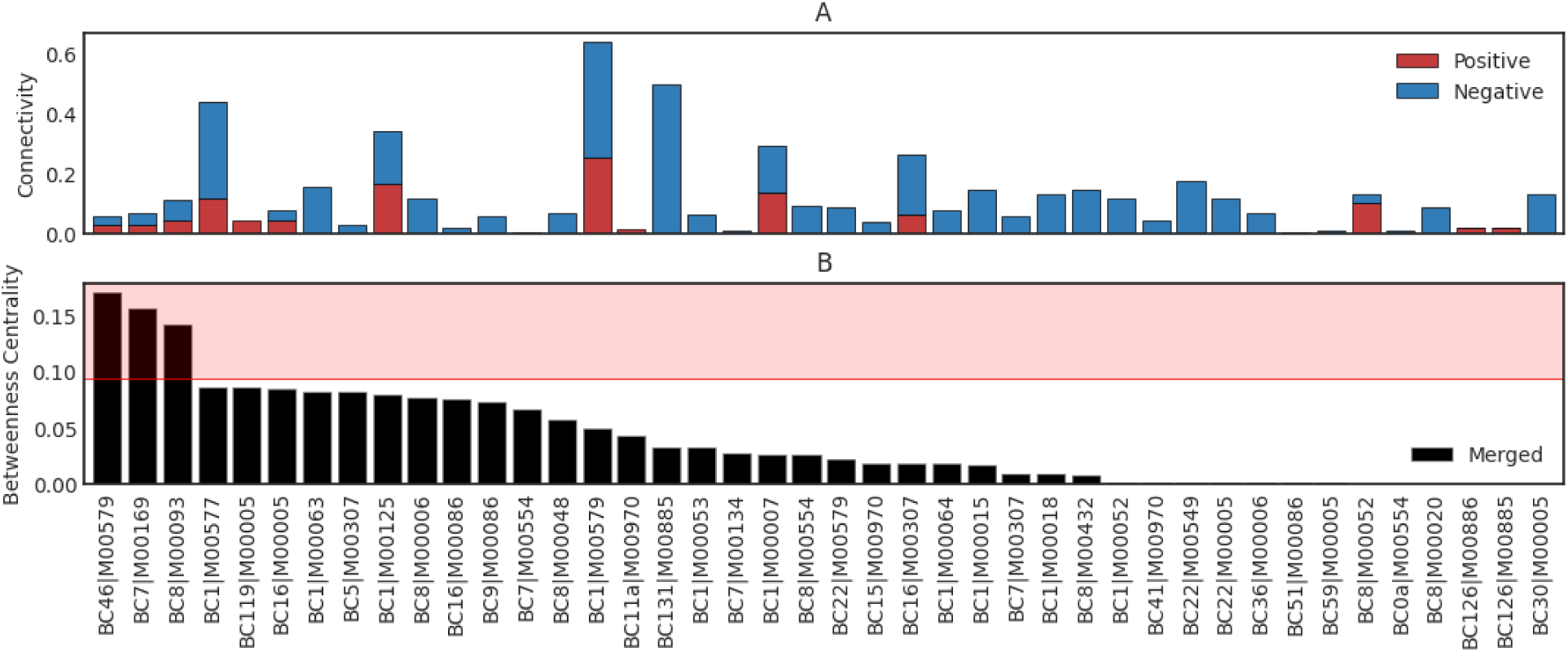
(A) Stacked barchart of *BiDirectional Clustered Networks* connectivities enriched in caries cohort (red) and caries-free cohort (blue). (B) Barchart of betweenness centrality for the union of the caries-enriched and caries-free-enriched networks with outlier bridge nodes identified using a robust outlier threshold (median + 3 MAD) of betweenness centrality.

Analysis of betweenness-centrality revealed three distinct outliers with disproportionately high values: 1) *Granulicatella adiacens* phosphate acetyltransferase-acetate kinase conversion from acetyl-CoA to acetate (*BC46|M00579,* value=0.17); 2) *Fusobacterium polymorphum* CAM pathway (*BC7|M00169,* value=0.16); and 3) *Haemophilus_D parainfluenzae* phosphatidylethanolamine biosynthesis (*BC8|M00093,* value=0.13) (Fig. 6, Table S5).

The *BC7|M00169* instance brings up an important caveat regarding false positives in *KEGG Brite Hierarchy,* that is, *Fusobacterium polymorphum* is an anaerobic Gram-negative bacterium that relies on fermentation for energy in oxygen-free environments (e.g., human oral cavity) and CAM is a plant-specific carbon fixation pathway. However, there is a substantial overlap in *KEGG* orthologs between M00169 and several other *KEGG* modules including M00172 C4-dicarboxylic acid cycle (100% overlap coefficient) as well as M00171 C4-dicarboxylic acid cycle and M00173 reductive citrate cycle (50% overlap coefficient) as shown in Table S6 and documented previously (45). Thus, the source of these false positive assignments result from HMM score thresholds for *KEGG* orthology and are not an issue with *Leviathan* or the network-based analysis methods.

These three bridge TRFM appeared in both caries-enriched and caries-free-enriched subnetworks but with distinct co-expression partners. In the caries-free-enriched subnetwork, *G. adiacens M00579* co-expressed with FAS II fatty acid elongation modules in *Capnocytophaga sputigena*, *Streptococcus* spp., and *Streptococcus sanguinis*; *F. polymorphum* M00169 coexpressed with *Streptococcus* FAS II modules and its own histidine degradation and pyruvate oxidation pathways; and *H. parainfluenzae* M00093 co-expressed predominantly with its own central carbon metabolism (gluconeogenesis, pentose phosphate pathway, fumarate reductase) and *Streptococcus* glycolysis and glycogen biosynthesis. In the caries-enriched subnetwork, all three bridge nodes shifted toward inter-species coexpression with *Veillonella parvula* cofactor biosynthesis (biotin, riboflavin), *Streptococcus* branched-chain amino acid biosynthesis, and *F. polymorphum* glycogen biosynthesis, consistent with cross-feeding patterns associated with cariogenic biofilms in the ecological caries hypothesis (46).

This case study demonstrates that *Leviathan’s* pathway profiling, when paired with differential co-expression network analysis, can resolve organism-specific transcriptional differences between health and disease ecological states that would be undetectable with taxon-level or pathway-level profiling alone.

## Discussion

We introduced *Leviathan*, a modular software package for integrated taxonomic and functional profiling of metagenomes and metatranscriptomes against user-defined (pan)genome-resolved reference catalogs. *Leviathan* is designed for both taxonomic and functional profiling by leveraging the strengths of ultrafast methods such as *Sylph* (1) and *Salmon* (23) while also building on prior conceptual work pioneered by *HUMAnN* (6). The benchmarking reveals that *Leviathan’s* pseudo-alignment backend retains competitive alignment classification performance while improving computational efficiency of functional profiling when compared to traditional alignment or translated search methods.

*Leviathan* is designed for the increasing need for support of user-defined reference databases. In this paradigm, the reference is a parameter and comparing *Leviathan* end-to-end functional profiling against profilers that use different databases would conflate database composition, annotation scheme, and post-alignment processing with alignment strategy. To address the read assignment bottlenecks, we held the reference constant by using a single CDS catalog derived from each CAMI dataset’s source genomes and varied only the read-assignment backend and its post-filtering. This isolates the step at which the DNA-space and protein-space strategies diverge, and which dominates runtime in translated-search designs. In the context of *HUMAnN*, this strategy quantifies two ends of the alignment spectrum, that is, if all reads were assigned in DNA-space or protein-space.

With our benchmarking strategy, we found that backend choice, either pseudo-alignment or traditional alignment, within DNA-space has little effect on classification accuracy. However, with comparable accuracy and equivalent peak memory usage pseudo-alignment methods had a 2.1 to 3.4 fold lower runtime. All DNA-space backends were evaluated on primary alignments to provide a comparable evaluation unit, as each resolves multi-mapping through distinct downstream mechanisms (expectation maximization, identity thresholds, or gene coverage filters) that are not directly comparable at the alignment level. Second, the majority of genome-level error is within-species across the three DNA-space backends as aggregation to ANI-defined genome clusters recovered accuracy from ∼90% at the genome level to ∼97% at the pangenome level demonstrating the robustness of the pangenome as the ideal unit for functional analysis in complex communities. Third, translated search produces a different error profile rather than simply a lower accuracy. Each DNA-space backend assigns every read to a single genome, so micro-averaged precision, recall, and accuracy are identical for all three. *HUMAnN’s Diamond* translated search configuration retains, for each read, every alignment scoring within one percent of that read’s best hit, so a single read can be assigned to several genomes at once. Because synonymous nucleotide variation collapses in protein-space, reads from conserved coding regions become compatible with proteins from several source genomes, and each additional incorrect hit contributes a false positive without a corresponding false negative. Precision therefore falls sharply while recall is largely preserved, giving a median genome-level precision of 34.2% against a recall of 78.6%. However, this design is intentional and a strength for *HUMAnN* as the standard workflow involves DNA-space alignments as a first pass and where unplaced reads for remote homology can be searched in protein-space alignments which is out of scope for *Leviathan*.

These results do not establish that DNA-space search is superior to protein-space search. The two strategies trade against one another. Translated search tolerates nucleotide divergence and can recover conserved gene families from organisms with no close nucleotide representative in the catalog, which is why *HUMAnN* employs it as a fallback tier and why it remains the appropriate choice when the reference is incomplete. *Leviathan* is scoped accordingly: it quantifies a community against a reference the user defines, and organisms absent from that reference are not profiled. Two mechanisms enforce this boundary. *Leviathan* applies a stringent alignment score threshold by default, so a read that cannot align at high identity to a reference CDS is left unassigned rather than being attributed to the closest available match. Separately, *Sylph* reports the estimated fraction of the community not represented in the reference so unprofiled diversity is directly estimated.

Cross-catalog benchmarking against *OceanDNA* species representatives confirms that these boundaries hold in practice: reads from represented genomes achieved 94.2% assignment precision while reads from unrepresented genomes showed a 0.2% false-positive rate (Fig. S2), indicating that the stringent alignment threshold suppresses spurious assignments without sacrificing precision for organisms present in the catalog.

While individual members of a pangenome can carry identical coding sequences, a read mapping to such a gene could have originated from any of those genomes and assigning it to one is inherently ambiguous. Aggregation to the pangenome level removes this ambiguity, which accounts for most genome-level error and is why all three DNA-space backends recover accuracy at the pangenome level. Our reference retained every sample-specific strain, maximizing within-species redundancy and therefore constituting the most adverse case for read assignment; the genome-level accuracies are accordingly a lower bound. *Leviathan* places no constraint on the composition of the reference and pangenome definitions are optional, so dereplicated collections are equally valid inputs and no aggregation layer is required when each genome already represents a distinct species cluster. Our benchmarks and case studies used the full multi-strain catalog rather than a dereplicated subset because selecting a single representative per species, while more computationally efficient for large catalogs, discards accessory genes carried by other members, and it is precisely these genes that distinguish strains and contribute to functional diversity within a pangenome. Retaining the union of strains ensures that the full gene repertoire of each pangenome is available for quantification, which is the central motivation for pangenome-resolved profiling. In practice, we recommend users choose between these strategies based on their analytical priorities: pangenome-resolved catalogs that retain all constituent genomes capture the full functional repertoire of a species cluster at the cost of higher resource usage, while dereplicated catalogs that retain a single representative per species reduce computational requirements but may not cover accessory genes present in other members of the pangenome.

The practical difficulty with existing tools is obtaining their full functionality when the reference is user-defined. *HUMAnN’s* pathway inference is reached transitively: custom proteins must be aligned to *UniRef* clusters, the functional annotation assigned to each cluster is then mapped to *MetaCyc* reactions, and reactions to pathways via *MinPath*. The protein alignment approach uses a single global identity threshold rather than marker-specific cutoffs as with HMMs and the functional annotation a read ultimately receives is determined by the *UniRef* cluster it matches rather than by direct evaluation against a curated model. Further, each transitive mapping step can also lose coverage for annotations that lack cross-references between databases and could underestimate the pathway coverage. *Leviathan’s* functional profiling logic is agnostic to annotation methodology, supporting both HMM and alignment-based, but the standard workflow recommends annotating translated coding sequences directly using HMMs with hits filtered by curated or trusted score thresholds so annotations are direct rather than inherited. Though, this trade runs in both directions: curated thresholds raise annotation precision but a divergent coding sequence that a transitive mapping would annotate by remote homology may receive no feature assignment at all.

*Leviathan* is designed for custom reference catalogs, but assembling, binning, and annotating genomes from metagenomic data can be computationally expensive. These costs are not unique to *Leviathan*, as assembly, binning, and annotation are standard steps in any genome-resolved metagenomics workflow. The *Leviathan* index itself is built once from the resulting catalog and reused across every sample in a cohort. For datasets where genome-resolved approaches are not feasible, reference-based profiling methods such as *HUMAnN* or *Meteor2* are recommended over targeted methods like *Leviathan*.

*Leviathan* has several limitations. For instance, it is not a substitute for translated search in reference-poor settings and users should verify reference coverage per dataset using *Sylph’s* estimated unknown fraction. Currently, only paired-end and single-end short reads are supported but long-reads will be supported with *Oarfish* (47) when *CAMI* provides a synthetic metatranscriptomic dataset to benchmark. Pathway abundance is the sum of all feature abundances within that pathway (e.g., *KEGG* orthologs) for a given genomic unit and a feature present in multiple modules contributes its full abundance to each rather than being divided across them. This is a deliberate design choice: pathways are an overlapping cover of feature space, not a partition, and an enzyme that participates in both glycolysis and pyruvate metabolism is genuine evidence for both. Dividing by the number of modules would make each pathway’s abundance depend on how many other modules happen to share its orthologs, which is an artifact of *KEGG’s* curation density rather than a property of the community. Two consequences follow: 1) pathway abundance scales with the number of steps in a pathway and is not size-normalized; 2), modules with high ortholog overlap receive similar or identical abundances by construction. The M00169 and M00172 pair documented in *Case Study II* has an overlap coefficient of 1.0 and cannot be distinguished by pathway abundance at all. Pathway abundance is therefore intended for comparing the same pathway across samples or genomic units, while pathway coverage, which is bounded in [0, 1] and size-normalized, is the appropriate metric for comparing across pathways within a sample. The module overlap coefficients in Table S6 are a necessary interpretive companion to any abundance table, as they identify which pathway pairs are confounded by shared orthologs. Another limitation is that full functional support partially relies on *KEGG*, but the modular architecture is designed to readily incorporate other functional databases such as *MetaCyc* if pathway graphs are available which would broaden its utility. Future work will focus on expanding this database support and further optimizing data structures for even larger-scale meta-analyses.

A key advance offered by *Leviathan* is its seamless, native support for pangenome-resolved analysis and flexibility for custom reference databases. *Leviathan* streamlines this entirely, integrating pangenome definitions at the indexing stage and automatically generating both genome and pangenome level outputs. This functionality makes it trivial for researchers to investigate critical biological questions related to functional redundancy, niche specialization, and the distribution of metabolic capabilities across related microbial populations.

In conclusion, functional profilers such as *Leviathan*, *HUMAnN*, *Meteor2*, *Woltka*, and *MIDAS* occupy complementary positions in a shared design space, and many studies will benefit from both a reference-free and a reference-resolved view of the same data. *Leviathan*, in particular, makes (pan)genome-resolved functional profiling against user-defined references practical at cohort scale, paving the way for new discoveries in microbial ecology.

## Data Availability

*Leviathan* is an open-sourced software package available at https://github.com/jolespin/leviathan. *CAMI* datasets were downloaded from *GigaDB* (https://gigadb.org/dataset/100344). *CAMI* benchmarking and intermediate files are available on Zenodo (doi: 10.5281/zenodo.20172866). The *Marine Plastisphere Microbiome* case study metagenomes were downloaded from NCBI BioProject PRJNA777294 and source code is available at https://github.com/jolespin/leviathan-case-study-plastisphere. The *Dental Caries Oral Microbiome* case study metatranscriptomes were downloaded from NCBI BioProject PRJNA383868 source code is available at https://github.com/jolespin/leviathan-case-study-oral.

## Acknowledgements

*HUMAnN* developers for laying the foundation for this research. The founders of *NewAtlantis Labs* for their support in the development of this software for marine conservation. R01AI170111 to CLD.

## Supplemental Materials

**Figure S1.**
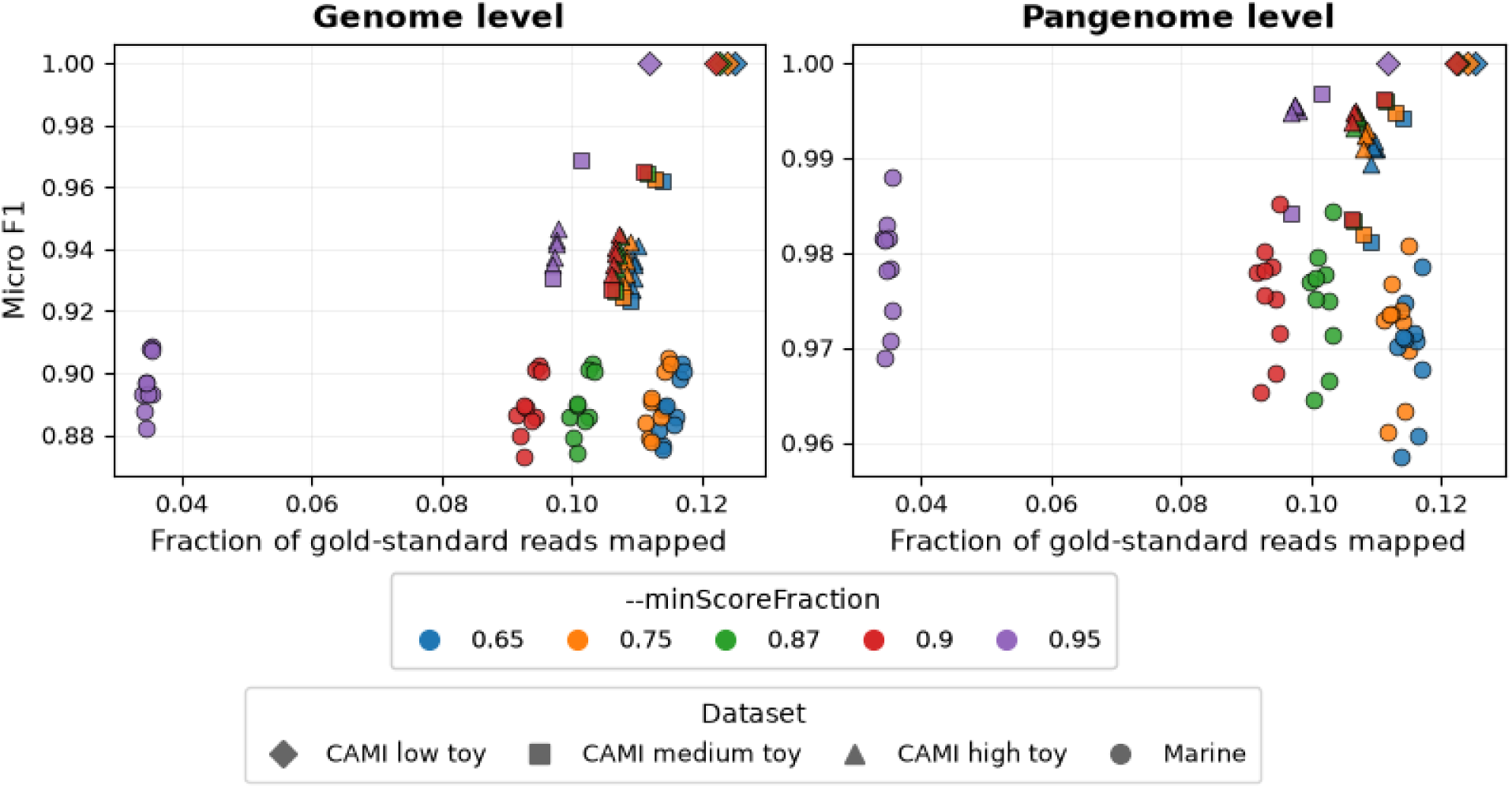
*Salmon --minScoreFraction* parameter sweep. Micro F1 versus the fraction of gold-standard reads mapped to CDS for five *--minScoreFraction* values (0.65 - 0.95) across all *CAMI* benchmark datasets at genome (left) and pangenome (right) levels. Each point represents one sample.

**Figure S2.**
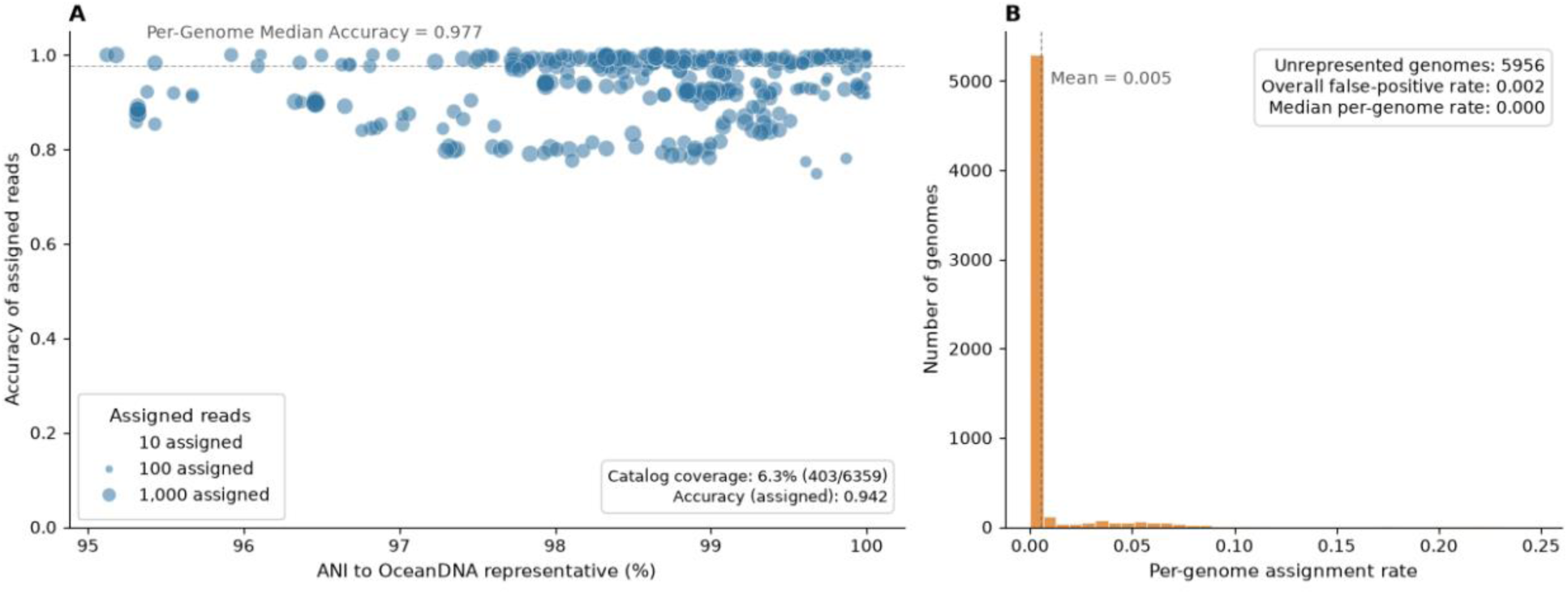
Cross-catalog benchmarking of *CAMI-II* Marine reads profiled against *OceanDNA* species-level representatives using *Salmon* with *Leviathan’s* default configuration. (A) Per-genome accuracy of read assignment for represented genomes (ANI ≥ 95%, alignment fraction ≥ 50%) versus ANI to the matched *OceanDNA* representative. Marker size is proportional to the number of assigned reads. Dashed line indicates the per-genome median accuracy (0.977); (B) Distribution of per-genome assignment rates for unrepresented genomes (n = 5,956), showing the fraction of reads from each genome that received any *OceanDNA* assignment.

**Figure S3** - Interactive network version of *BiDirectional Clustered Network* with metadata hover tool

**Table S1** - Benchmarking summary statistics. Per-sample runtime, peak memory usage, and classification performance at genome and pangenome levels for each alignment backend across *CAMI-I* and *CAMI-II* datasets, *Leviathan* end-to-end profiling statistics for benchmarking and case study datasets, cross-catalog benchmarking metrics for *CAMI-II* Marine reads profiled against *OceanDNA* species representatives, *Salmon --minScoreFraction* parameter sweep benchmarking, and *Skani* ANI calculations for *CAMI-II* Marine against *OceanDNA*.

**Table S2** - Differential abundance of pangenome-level taxonomic profiling via *ANCOM-BC* of mature vs. early stage plastic biofilms.

**Table S3** - Differential coverage of pangenome-level pathway profiling via Mann Whitney-U test and rank-biserial correlation of mature vs. early stage plastic biofilms.

**Table S4** - *BiDirectional Clustered Network* edge weights and annotations for caries-enriched and caries-free-enriched *Differential Ensemble Co-expression Networks*

**Table S5** - *BiDirectional Clustered Networks* Leiden communities, connectivities, and betweenness-centrality node-level statistics

**Table S6** - Overlap coefficients between *KEGG* modules based *KEGG* ortholog sets

**Text S1** - Specific details on commands used for benchmarking

## Notes

https://github.com/jolespin/leviathan

### Competing Interest Statement

Josh Espinoza developed this open-source software while employed at NewAtlantis Labs

### Summary of Updates

The most significant changes are summarized below: Benchmark reframe: Rather than presenting a tool-versus-tool comparison between Leviathan and HUMAnN, we now benchmark four read-assignment backends against a common CDS catalog derived from each CAMI dataset's reference genomes: (1) Salmon pseudo-alignment using Leviathan's configuration, (2) Bowtie2 alignment using HUMAnN's configuration, (3) Bowtie2 alignment using Meteor2's configuration, and (4) Diamond blastx translated search using HUMAnN's configuration. This design isolates the core read-assignment bottleneck backend step from confounding variables such as database composition, annotation scheme, and post-alignment processing. All backend parameters are documented in Supplemental Text S1. Repositioned scope: The Introduction now explicitly distinguishes two reference paradigms, fixed reference catalogs and study-specific reference catalogs, and states that neither is inherently more legitimate while providing pros and cons of both approaches. The Discussion recommends existing reference-based profilers such as HUMAnN or Meteor2 for datasets where genome-resolved approaches are not feasible. Cross-catalog benchmarking: We added benchmarking against OceanDNA species representatives, an independent pre-compiled collection of 52,325 genomes not derived from the CAMI source genomes. This directly addresses concerns about false-positive behavior when the reference is independent of the study: reads from represented genomes achieved 94.2% assignment accuracy while reads from unrepresented genomes showed a 0.2% false-positive rate. Comprehensive classification metrics: All “up to XX%” claims have been replaced with medians. We now report precision, recall, and F1 at both genome and pangenome levels (new Figure 2), a Salmon --minScoreFraction parameter sweep (Figure S1), and complete benchmarking statistics (Table S1). Corrected HUMAnN characterization: The manuscript now accurately describes HUMAnN's two-tier strategy (DNA-space first pass, protein-space fallback) and corrects the read-handling description from merging to concatenation. Expanded methods: We added detailed descriptions of Leviathan’s preprocessing module, Taxa-Resolved Functional Module computation, module independence between Sylph and Salmon, and pathway abundance interpretation, along with an expanded Limitations section.

## References

1. Shaw J, Yu YW. 2024. Rapid species-level metagenome profiling and containment estimation with sylph. Nat Biotechnol 1–12.

2. Wood DE, Lu J, Langmead B. 2019. Improved metagenomic analysis with Kraken 2. Genome Biol 20:257.

3. Piro VC, Dadi TH, Seiler E, Reinert K, Renard BY. 2020. ganon: precise metagenomics classification against large and up-to-date sets of reference sequences. Bioinformatics 36:i12–i20.

4. Blanco-Míguez A, Beghini F, Cumbo F, McIver LJ, Thompson KN, Zolfo M, Manghi P, Dubois L, Huang KD, Thomas AM, Nickols WA, Piccinno G, Piperni E, Punčochář M, Valles-Colomer M, Tett A, Giordano F, Davies R, Wolf J, Berry SE, Spector TD, Franzosa EA, Pasolli E, Asnicar F, Huttenhower C, Segata N. 2023. Extending and improving metagenomic taxonomic profiling with uncharacterized species using MetaPhlAn 4. Nat Biotechnol 41:1633–1644.

5. Ruscheweyh H-J, Milanese A, Paoli L, Sintsova A, Mende DR, Zeller G, Sunagawa S. 2021. mOTUs: Profiling Taxonomic Composition, Transcriptional Activity and Strain Populations of Microbial Communities. Curr Protoc 1:e218.

6. Beghini F, McIver LJ, Blanco-Míguez A, Dubois L, Asnicar F, Maharjan S, Mailyan A, Manghi P, Scholz M, Thomas AM, Valles-Colomer M, Weingart G, Zhang Y, Zolfo M, Huttenhower C, Franzosa EA, Segata N. 2021. Integrating taxonomic, functional, and strain-level profiling of diverse microbial communities with bioBakery 3. eLife 10:e65088.

7. Ghozlane A, Thirion F, Plaza Oñate F, Gauthier F, Le Chatelier E, Annamalé A, Almeida M, Ehrlich SD, Pons N. 2025. Accurate profiling of microbial communities for shotgun metagenomic sequencing with Meteor2. Microbiome 13:227.

8. Speth DR, Pullen N, Aroney STN, Coltman BL, Osvatic J, Woodcroft BJ, Rattei T, Wagner M. 2025. GlobDB: a comprehensive species-dereplicated microbial genome resource. Bioinforma Adv 5:vbaf280.

9. Parks DH, Chuvochina M, Rinke C, Mussig AJ, Chaumeil P-A, Hugenholtz P. 2022. GTDB: an ongoing census of bacterial and archaeal diversity through a phylogenetically consistent, rank normalized and complete genome-based taxonomy. Nucleic Acids Res 50:D785–D794.

10. Nishimura Y, Yoshizawa S. 2022. The OceanDNA MAG catalog contains over 50,000 prokaryotic genomes originated from various marine environments. Sci Data 9:305.

11. Ma B, Lu C, Wang Y, Yu J, Zhao K, Xue R, Ren H, Lv X, Pan R, Zhang J, Zhu Y, Xu J. 2023. A genomic catalogue of soil microbiomes boosts mining of biodiversity and genetic resources. Nat Commun 14:7318.

12. Espinoza JL, Phillips A, Prentice MB, Tan GS, Kamath PL, Lloyd KG, Dupont CL. 2024. Unveiling the microbial realm with VEBA 2.0: a modular bioinformatics suite for end-to-end genome-resolved prokaryotic, (micro)eukaryotic and viral multi-omics from either short- or long-read sequencing. Nucleic Acids Res 52:e63.

13. Espinoza JL, Dupont CL. 2022. VEBA: a modular end-to-end suite for in silico recovery, clustering, and analysis of prokaryotic, microeukaryotic, and viral genomes from metagenomes. BMC Bioinformatics 23:419.

14. Shaiber A, Eren AM. 2019. Composite Metagenome-Assembled Genomes Reduce the Quality of Public Genome Repositories. mBio 10:10.1128/mbio.00725-19.

15. Olm MR, Brown CT, Brooks B, Banfield JF. 2017. dRep: a tool for fast and accurate genomic comparisons that enables improved genome recovery from metagenomes through de-replication. ISME J 11:2864–2868.

16. Shaw J, Yu YW. 2023. Fast and robust metagenomic sequence comparison through sparse chaining with skani. Nat Methods 20:1661–1665.

17. Molina-Pardines C, Haro-Moreno JM, Rodriguez-Valera F, López-Pérez M. 2025. Extensive paralogism in the environmental pangenome: a key factor in the ecological success of natural SAR11 populations. Microbiome 13:41.

18. Zhu Q, Huang S, Gonzalez A, McGrath I, McDonald D, Haiminen N, Armstrong G, Vázquez-Baeza Y, Yu J, Kuczynski J, Sepich-Poore GD, Swafford AD, Das P, Shaffer JP, Lejzerowicz F, Belda-Ferre P, Havulinna AS, Méric G, Niiranen T, Lahti L, Salomaa V, Kim H-C, Jain M, Inouye M, Gilbert JA, Knight R. 2022. Phylogeny-Aware Analysis of Metagenome Community Ecology Based on Matched Reference Genomes while Bypassing Taxonomy. mSystems 7:e00167–22.

19. Zhao C, Dimitrov B, Goldman M, Nayfach S, Pollard KS. 2023. MIDAS2: Metagenomic Intra-species Diversity Analysis System. Bioinformatics 39:btac713.

20. Sczyrba A, Hofmann P, Belmann P, Koslicki D, Janssen S, Dröge J, Gregor I, Majda S, Fiedler J, Dahms E, Bremges A, Fritz A, Garrido-Oter R, Jørgensen TS, Shapiro N, Blood PD, Gurevich A, Bai Y, Turaev D, DeMaere MZ, Chikhi R, Nagarajan N, Quince C, Meyer F, Balvočiūtė M, Hansen LH, Sørensen SJ, Chia BKH, Denis B, Froula JL, Wang Z, Egan R, Don Kang D, Cook JJ, Deltel C, Beckstette M, Lemaitre C, Peterlongo P, Rizk G, Lavenier D, Wu Y-W, Singer SW, Jain C, Strous M, Klingenberg H, Meinicke P, Barton MD, Lingner T, Lin H-H, Liao Y-C, Silva GGZ, Cuevas DA, Edwards RA, Saha S, Piro VC, Renard BY, Pop M, Klenk H-P, Göker M, Kyrpides NC, Woyke T, Vorholt JA, Schulze-Lefert P, Rubin EM, Darling AE, Rattei T, McHardy AC. 2017. Critical Assessment of Metagenome Interpretation—a benchmark of metagenomics software. Nat Methods 14:1063–1071.

21. Meyer F, Fritz A, Deng Z-L, Koslicki D, Lesker TR, Gurevich A, Robertson G, Alser M, Antipov D, Beghini F, Bertrand D, Brito JJ, Brown CT, Buchmann J, Buluç A, Chen B, Chikhi R, Clausen PTLC, Cristian A, Dabrowski PW, Darling AE, Egan R, Eskin E, Georganas E, Goltsman E, Gray MA, Hansen LH, Hofmeyr S, Huang P, Irber L, Jia H, Jørgensen TS, Kieser SD, Klemetsen T, Kola A, Kolmogorov M, Korobeynikov A, Kwan J, LaPierre N, Lemaitre C, Li C, Limasset A, Malcher-Miranda F, Mangul S, Marcelino VR, Marchet C, Marijon P, Meleshko D, Mende DR, Milanese A, Nagarajan N, Nissen J, Nurk S, Oliker L, Paoli L, Peterlongo P, Piro VC, Porter JS, Rasmussen S, Rees ER, Reinert K, Renard B, Robertsen EM, Rosen GL, Ruscheweyh H-J, Sarwal V, Segata N, Seiler E, Shi L, Sun F, Sunagawa S, Sørensen SJ, Thomas A, Tong C, Trajkovski M, Tremblay J, Uritskiy G, Vicedomini R, Wang Z, Wang Z, Wang Z, Warren A, Willassen NP, Yelick K, You R, Zeller G, Zhao Z, Zhu S, Zhu J, Garrido-Oter R, Gastmeier P, Hacquard S, Häußler S, Khaledi A, Maechler F, Mesny F, Radutoiu S, Schulze-Lefert P, Smit N, Strowig T, Bremges A, Sczyrba A, McHardy AC. 2022. Critical Assessment of Metagenome Interpretation: the second round of challenges. Nat Methods 19:429–440.

22. Bushnell B. 2014. BBMap: A Fast, Accurate, Splice-Aware Aligner. LBNL-7065E. Lawrence Berkeley National Lab. (LBNL), Berkeley, CA (United States).

23. Patro R, Duggal G, Love MI, Irizarry RA, Kingsford C. 2017. Salmon provides fast and bias-aware quantification of transcript expression. Nat Methods 14:417–419.

24. Aramaki T, Blanc-Mathieu R, Endo H, Ohkubo K, Kanehisa M, Goto S, Ogata H. 2020. KofamKOALA: KEGG Ortholog assignment based on profile HMM and adaptive score threshold. Bioinformatics 36:2251–2252.

25. Srivastava A, Malik L, Smith T, Sudbery I, Patro R. 2019. Alevin efficiently estimates accurate gene abundances from dscRNA-seq data. Genome Biol 20:65.

26. Hoyer S, Hamman J. 2017. xarray: N-D labeled Arrays and Datasets in Python. J Open Res Softw 5.

27. Finn RD, Clements J, Eddy SR. 2011. HMMER web server: interactive sequence similarity searching. Nucleic Acids Res 39:W29–W37.

28. Larralde M, Zeller G. 2023. PyHMMER: a Python library binding to HMMER for efficient sequence analysis. Bioinformatics 39:btad214.

29. Richardson L, Allen B, Baldi G, Beracochea M, Bileschi ML, Burdett T, Burgin J, Caballero-Pérez J, Cochrane G, Colwell LJ, Curtis T, Escobar-Zepeda A, Gurbich TA, Kale V, Korobeynikov A, Raj S, Rogers AB, Sakharova E, Sanchez S, Wilkinson DJ, Finn RD. 2023. MGnify: the microbiome sequence data analysis resource in 2023. Nucleic Acids Res 51:D753–D759.

30. Aton M, McDonald D, Cañardo Alastuey J, Azom R, Batra P, Bezshapkin V, Bolyen E, Cagle A, Caporaso JG, Debelius JW, Gorlick K, Hamsanipally N, Hunger L, Keluskar A, Liao D, Lu YY, Navas-Molina JA, Pitman A, Rideout JR, Sazonov A, Sathappan B, Schwarzberg Lipson K, Sfiligoi I, Tapo C, Vázquez-Baeza Y, Wu Z, Xu ZZ, Ye MS, Zhao J, Knight R, Morton JT, Zhu Q. 2026. Scikit-bio: a fundamental Python library for biological omic data analysis. Nat Methods 23:274–276.

31. Lin H, Peddada SD. 2020. Analysis of compositions of microbiomes with bias correction. Nat Commun 11:3514.

32. Seabold S, Perktold J. 2010. Statsmodels: Econometric and Statistical Modeling with Python. SciPy 2010 10.25080/Majora-92bf1922-011.

33. Virtanen P, Gommers R, Oliphant TE, Haberland M, Reddy T, Cournapeau D, Burovski E, Peterson P, Weckesser W, Bright J, van der Walt SJ, Brett M, Wilson J, Millman KJ, Mayorov N, Nelson ARJ, Jones E, Kern R, Larson E, Carey CJ, Polat İ, Feng Y, Moore EW, VanderPlas J, Laxalde D, Perktold J, Cimrman R, Henriksen I, Quintero EA, Harris CR, Archibald AM, Ribeiro AH, Pedregosa F, van Mulbregt P. 2020. SciPy 1.0: fundamental algorithms for scientific computing in Python. Nat Methods 17:261–272.

34. Espinoza JL, Shah N, Singh S, Nelson KE, Dupont CL. 2020. Applications of weighted association networks applied to compositional data in biology. Environ Microbiol 22:3020–3038.

35. Espinoza JL, Torralba M, Leong P, Saffery R, Bockmann M, Kuelbs C, Singh S, Hughes T, Craig JM, Nelson KE, Dupont CL. 2022. Differential network analysis of oral microbiome metatranscriptomes identifies community scale metabolic restructuring in dental caries. PNAS Nexus 1:pgac239.

36. Traag VA, Waltman L, van Eck NJ. 2019. From Louvain to Leiden: guaranteeing well-connected communities. Sci Rep 9:5233.

37. Nabwera HM, Espinoza JL, Worwui A, Betts M, Okoi C, Sesay AK, Bancroft R, Agbla SC, Jarju S, Bradbury RS, Colley M, Jallow AT, Liu J, Houpt ER, Prentice AM, Antonio M, Bernstein RM, Dupont CL, Kwambana-Adams BA. 2021. Interactions between fecal gut microbiome, enteric pathogens, and energy regulating hormones among acutely malnourished rural Gambian children. eBioMedicine 73.

38. Salamzade R, Kottapalli A, Kalan LR. 2025. skDER and CiDDER: two scalable approaches for microbial genome dereplication. Microb Genomics 11:001438.

39. Chaumeil P-A, Mussig AJ, Hugenholtz P, Parks DH. 2020. GTDB-Tk: a toolkit to classify genomes with the Genome Taxonomy Database. Bioinformatics 36:1925–1927.

40. Quinn TP, Erb I. 2020. Amalgams: data-driven amalgamation for the dimensionality reduction of compositional data. NAR Genomics Bioinforma 2:lqaa076.

41. Suzek BE, Wang Y, Huang H, McGarvey PB, Wu CH, the UniProt Consortium. 2015. UniRef clusters: a comprehensive and scalable alternative for improving sequence similarity searches. Bioinformatics 31:926–932.

42. Bos RP, Kaul D, Zettler ER, Hoffman JM, Dupont CL, Amaral-Zettler LA, Mincer TJ. 2023. Plastics select for distinct early colonizing microbial populations with reproducible traits across environmental gradients. Environ Microbiol 25:2761–2775.

43. Espinoza JL, Harkins DM, Torralba M, Gomez A, Highlander SK, Jones MB, Leong P, Saffery R, Bockmann M, Kuelbs C, Inman JM, Hughes T, Craig JM, Nelson KE, Dupont CL. 2018. Supragingival Plaque Microbiome Ecology and Functional Potential in the Context of Health and Disease. mBio 9:e01631–18.

44. Robaina-Estévez S, Gutiérrez J. 2024. Applications of marine microbial community models in the nature-based economy. PLOS Sustain Transform 3:e0000145.

45. Darzi Y, Falony G, Vieira-Silva S, Raes J. 2016. Towards biome-specific analysis of meta-omics data. ISME J 10:1025–1028.

46. Takahashi N, Nyvad B. 2011. The role of bacteria in the caries process: ecological perspectives. J Dent Res 90:294–303.

47. Zare Jousheghani Z, Singh NP, Patro R. 2025. Oarfish: enhanced probabilistic modeling leads to improved accuracy in long read transcriptome quantification. Bioinformatics 41:i304–i313.

48. Sunagawa S, Acinas SG, Bork P, Bowler C, Eveillard D, Gorsky G, Guidi L, Iudicone D, Karsenti E, Lombard F, Ogata H, Pesant S, Sullivan MB, Wincker P, de Vargas C. 2020. Tara Oceans: towards global ocean ecosystems biology. Nat Rev Microbiol 18:428–445.

49. Rusch DB, Halpern AL, Sutton G, Heidelberg KB, Williamson S, Yooseph S, Wu D, Eisen JA, Hoffman JM, Remington K, Beeson K, Tran B, Smith H, Baden-Tillson H, Stewart C, Thorpe J, Freeman J, Andrews-Pfannkoch C, Venter JE, Li K, Kravitz S, Heidelberg JF, Utterback T, Rogers Y-H, Falcón LI, Souza V, Bonilla-Rosso G, Eguiarte LE, Karl DM, Sathyendranath S, Platt T, Bermingham E, Gallardo V, Tamayo-Castillo G, Ferrari MR, Strausberg RL, Nealson K, Friedman R, Frazier M, Venter JC. 2007. The Sorcerer II Global Ocean Sampling Expedition: Northwest Atlantic through Eastern Tropical Pacific. PLOS Biol 5:e77.

50. Caspi R, Billington R, Keseler IM, Kothari A, Krummenacker M, Midford PE, Ong WK, Paley S, Subhraveti P, Karp PD. 2020. The MetaCyc database of metabolic pathways and enzymes - a 2019 update. Nucleic Acids Res 48:D445–D453.

51. Ye Y, Doak TG. 2009. A Parsimony Approach to Biological Pathway Reconstruction/Inference for Genomes and Metagenomes. PLoS Comput Biol 5:e1000465.

