## Supplementary material for "Leviathan: A fast, memory-efficient, and scalable taxonomic and pathway profiler for (pan)genome-resolved metagenomics and metatranscriptomics": Text S1

#### Universal *CAMI* preprocessing

The following commands were run for low, medium, high, and marine complexity datasets separately:

##### Clustering genomes into pangenomes using *skani* and *VEBA*

skani triangle --sparse -t 16 -o ${out_dir}/skani_output.tsv -l ${genome_filepaths} --min-af 15 -s 80 -c 125 -m 1000 --ci

cat ${output_directory}/skani_output.tsv | cut -f1-5 | tail -n +2 | edgelist_to_clusters.py --basename -t 95 -a 50 --af_mode relaxed --cluster_prefix "PSLC-" -o ${output_directory}/genome_clusters.tsv --identifiers ${genome_identifiers} --export_graph ${output_directory}/networkx_graph.pkl --export_dict ${out_dir}/dict.pkl --export_representatives ${output_directory}/representatives.tsv

##### Predicting prokaryotic genes with *Pyrodigal*

pyrodigal -i ${fp} -a ${dir}/${id}.faa -d ${dir}/${id}.ffn -j 16 | append_geneid_to_prodigal_gff.py > ${dir}/${id}.gff

##### Quality assessing genomes with *CheckM2*

checkm2 predict -i ${genome_directory} -o ${output_directory} --threads 16 --genes -x faa --database_path ${checkm2_database}

### *Leviathan* Commands

##### Identifying *KEGG Ortholog* pathway markers:

pykofamsearch -i ${faa} -o ${out} -p=16 -b ${db}

##### Preprocessing genomes and annotations for *Leviathan*:

leviathan-preprocess.py -i ${manifest} -a ${annotations} -o ${output_directory} --annotation_format pykofamsearch

##### Building the *Leviathan* index:

leviathan-index.py -f ${fasta} -m ${feature_mapping} -g ${genomes} -d ${index_directory} -p=16 --pathway_database ${pathway_database}

##### Taxonomic profiling with *Leviathan*:

leviathan-profile-taxonomy.py -1 ${r1} -2 ${r2} -n ${id} -d ${index} -p=4 -o ${output_directory}

##### Functional profiling with *Leviathan*:

leviathan-profile-pathway.py -1 ${r1} -2 ${r2} -n ${id} -d ${index} -p=4 -o ${output_directory} --salmon_include_mappings --alignment_format sam

#### Building dataset-specific backend *CAMI* databases

##### Build *Salmon* index

salmon index --transcripts cds.fasta.gz --index salmon_index --keepDuplicates --threads 14

##### Build *Bowtie2* index

bowtie2-build --seed 0 -f cds.fasta.gz cds.fasta.gz --threads 14

##### Build *Diamond* database

```
# protein.fasta.gz has genes in the same order as cds.fasta.gz
diamond makedb --in protein.fasta.gz --db protein.dmnd --threads 14
```

#### *Leviathan* backend benchmarking against CAMI

##### Running *Salmon* *quant* backend with *Leviathan* config

salmon quant --skipQuant --meta --libType A --threads 4 --minScoreFraction 0.87 --index ${index} -1 ${r1} -2 ${r2} --output ${output_directory} --writeMappings ${output_directory}/mapped.sam

#### *HUMAnN* backend benchmarking against *CAMI*

##### Running *Bowtie2* backend with *HUMAnN* config

bowtie2 -q --no-unal --very-sensitive --no-hd --no-sq --threads 4 -U ${reads_concatenated} -x ${index} -S ${output_directory}/mapped.sam

##### Running *Diamond blastx* with *HUMAnN* config

diamond blastx --threads 4 --query {concatenated_reads_fasta} --evalue 1.0 --top 1 --sensitive --outfmt 6 --db ${db} --out ${output_directory}/diamond_results.tsv

#### *Meteor2* backend benchmarking against CAMI

##### Running *Bowtie2* backend with *Meteor2* config

bowtie2 --end-to-end --sensitive --trim-to 80 -k 100 --no-unal --threads 4 -U ${r1},${r2} -x ${index} -S ${output_directory}/mapped.sam
